# An autonomous meiosis-specific region of yeast Nup2 (hNup50) promotes normal meiotic chromosome dynamics in *Saccharomyces cerevisiae*

**DOI:** 10.1101/068056

**Authors:** Daniel B. Chu, Sean M. Burgess

## Abstract

Meiosis is a specialized cellular program required to create haploid gametes from diploid parent cells. Prior to the first meiotic division, homologous chromosomes pair, synapse, and recombine to ensure their proper disjunction at anaphase I. Additionally, telomeres tethered at the nuclear envelope cluster in the bouquet configuration where they are subjected to dramatic pulling forces acting from outside of the nucleus. In *Saccharomyces cerevisiae*, the telomere-associated protein Ndj1 is required for bouquet formation. When Ndj1 is absent, these dramatic motions cease and multiple steps of meiosis I prophase progression are delayed. Here we identified Nup2 in a pool of enriched proteins that co-purify with tagged Ndj1 from meiotic cell extracts. Nup2 is a nonessential nucleoporin that functions in nuclear transport, boundary activity, and telomere silencing in mitotically dividing cells. We found that deletion of *NUP2* delayed pairing and synapsis during meiosis, and led to decreased spore viability, similar to the *ndj1Δ* mutant phenotype. Surprisingly, the *nup2Δ ndj1Δ* double mutant failed to segregate chromosomes, even though the meiotic program continued. These results suggest that a physical impediment to nuclear division is created in the absence of Nup2 and Ndj1. Our deletion analysis of *NUP2* identified a previously uncharacterized 125-amino acid region that is both necessary and sufficient to complement all of *nup2Δ*’s meiotic phenotypes, which we call the <u>m</u>eiotic <u>a</u>utonomous <u>r</u>egion (MAR). We propose that Ndj1 and Nup2 function in parallel pathways to promote the dynamic chromosome events of meiotic chromosome dynamics, perhaps through the establishment or maintenance of higher-order chromosome organization.

## INTRODUCTION

Mutations affecting chromosome structure, organization, and recombination during meiosis I prophase often lead to segregation errors or the failure to execute the meiosis I division (MI; Zickler and Kleckner 2015). Chromosome segregation errors during meiosis are the leading cause of birth defects and developmental delays in humans (Hassold and Hunt 2001).

The events of meiotic prophase follow a specialized round of DNA replication when the meiotic chromosome axis is formed. The chromosome axis is composed of a linear array of loops of sister chromatids attached serially to an axial protein substrate (Zickler and Kleckner 1999; Kleckner 2006). This configuration directs nearly every chromosome-based event of meiotic prophase, including the regulation of double-strand break (DSB) formation, recombination partner choice (i.e. homolog versus sister chromatid), and acts as part of the meiotic checkpoint signaling apparatus (Kleckner *et al*. 2004). Over the course of meiotic prophase, the chromosome axes are aligned via Spo11-induced recombination interactions and ultimately joined along their lengths by the transverse element of the synaptonemal complex (SC), Zip1 (Sym *et al*. 1993; Keeney *et al*. 1997). Zip1 initially loads where DNA recombination intermediates have been stabilized during zygotene and proceeds to polymerize along the full length of chromosomes marking pachytene (Borner *et al*. 2004).

During the transition from leptotene to zygotene, telomeres attach to cytoskeletal components via a protein bridge composed of a trimer of SUN (Sad1/UNC-84) domain proteins, spanning the inner nuclear membrane, and a trimer of KASH (Klarsicht, ANC-1, Syne Homology) proteins that span the outer nuclear membrane (Sosa *et al*. 2012; Horn *et al*. 2013; Tapley and Starr 2013). In budding yeast, this configuration requires the telomere-associated protein, Ndj1, the SUN protein Mps3, and the KASH protein Csm4, which connects (directly or indirectly) to cytoskeletal actin (Figure 1A; Conrad *et al*. 1997; Conrad *et al*. 2007; Conrad *et al*. 2008). During zygotene, telomeres cluster near the spindle-pole body (SPB); during pachytene, telomeres disperse, yet remain attached to the nuclear envelope and are subject to actin-dependent pulling forces (Zickler and Kleckner 1998). In other organisms, actin and/or microtubules serve to direct telomere clustering and motion (Chikashige *et al*. 1994; Koszul and Kleckner 2009; Sheehan and Pawlowski 2009; Horn *et al*. 2013). Little is known about the role of the bouquet in meiosis, yet mutations that disrupt the bouquet also have defects in homolog pairing and recombination (Koszul and Kleckner 2009; Zickler and Kleckner 2016).

**Figure 1.**
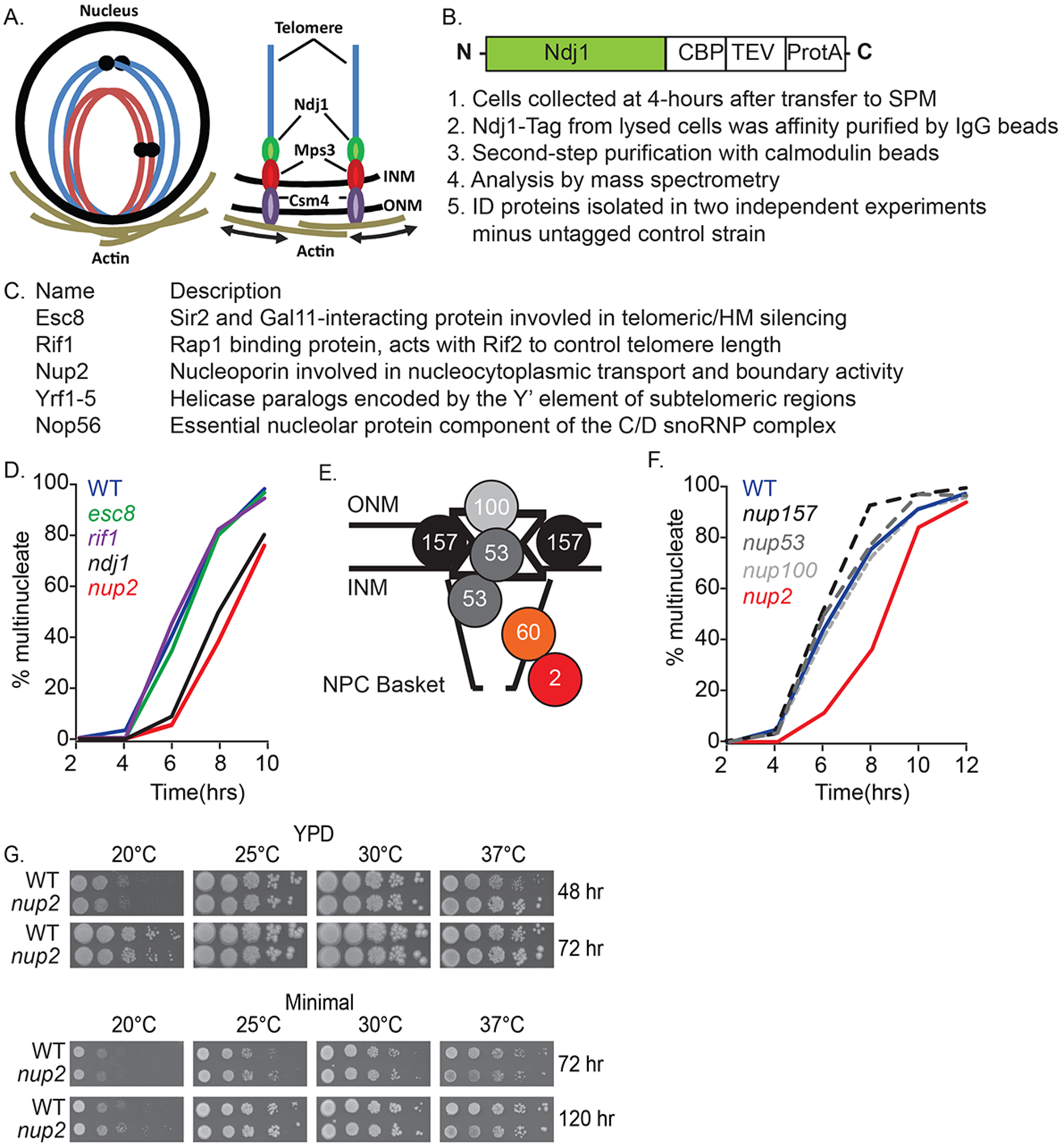
Identification of Nup2 and its role in meiotic progression. **A**. Schematic of the bouquet configuration of chromosomes during meiotic prophase I. The spatial arrangement of chromosomes with telomeres clustered and attached at the inner nuclear membrane is shown. These attachment sites are linked to actin-bundles that surround the nucleus via an Ndj1-Mps3-Csm4 protein bridge that spans the inner and outer nuclear membranes. The arrows depict the actin-directed motion that occurs through this linkage. **B**. Schematic of the C-terminal tagged Ndj1 protein for tandem-affinity purification (TAP) and an overview of the purification and analysis pipeline. CPB is the calmodulin-binding peptide, TEV is the TEV protease cleavage site and ProtA is the IgG binding domain of *Staphylococcus aureus* protein A. **C**. Proteins enriched after two-step purification of Ndj1 isolated from lysed cells at mid-meiotic prophase I with the exception of protein components of the ribosome (11), spliceosome (Prp31) and cytoplasmic proteins (Pet9, and Prr1; File S1). **D**. Kinetics of nuclear division in a time-course experiment from synchronized cells cultured in liquid sporulation medium (SPM). Analyzed cells were removed from the culture at the indicated hours after their initial transfer from an overnight YPA culture. At least 200 cells were analyzed for the presence of one, or more than one, DAPI-staining body. The percent of spore viability for each strain is indicated. All strains are diploid and are isogenic to the WT strain WT (Blue; SBY1903) except for *esc8Δ* (Green; SBY3942), *rif1Δ* (Purple; SBY3948), *nup2Δ* (Red; SBY3945), and *ndj1Δ* (Black; SBY1904). The kinetics for nuclear division of *rif2Δ and* the *rif1Δ rif2Δ* double mutant are shown in Figure S1. **E**. Schematic of the NPC with the relative positions of proteins analyzed in this study, including the outer-nuclear membrane (ONM), the inner-nuclear membrane (INM), the central channel, and the nucleoplasmic NPC basket. **F**. Kinetics of nuclear division from a time course experiment as described in part A. WT (Blue; SBY1903), *nup157Δ* (Black dashed; SBY5256), *nup53Δ* (Dark gray dashed; SBY5437), *nup100Δ* (Light gray dashed; SBY5549), and *nup2Δ* (Red; SBY4102). **G**. Lack of growth phenotype for the *nup2Δ* mutant under different incubation temperatures on rich (YPD) and minimal media. WT is SBY SBY4102). The growth phenotypes of all mutants are shown in Figure S2.

Telomere-led, actin-driven chromosome movements serve to improve the efficiency of pairing and recombination and may act to take apart nonspecific chromosome interactions and/or resolve interlocks that form during synapsis (Koszul *et al*. 2008; Koszul and Kleckner 2009; Lee *et al*. 2012; Lui *et al*. 2013). Loss of *NDJ1* results in defective meiotic chromosome dynamics leading to reduced sporulation efficiency and spore viability, as well as delayed meiotic progression due to activation of the recombination checkpoint (Chua and Roeder 1997; Conrad *et al*. 1997; Rockmill and Roeder 1998; Wu and Burgess 2006). Deleting *SPO11* bypasses checkpoint-induced delay or arrest conferred by a number of mutations affecting meiotic chromosome dynamics, including *ndj1Δ and* other mutations affecting the bouquet and chromosome motion (Alani *et al*. 1990; Cao *et al*. 1990; Bishop *et al*. 1992; Wu and Burgess 2006; Kosaka *et al*. 2008; Wanat *et al*. 2008).

Another integral feature of the nuclear envelope during meiotic prophase is the duplication of the SPB. The SPB is the microtubule organizing center in budding yeast and is required for chromosome separation in meiosis and mitosis (Lim *et al*. 2009). During meiosis, SPBs duplicate in coordination with DNA replication (Moens and Rapport 1971) but do not separate until the end of pachytene when the Ndt80 transcription factor is activated (Xu *et al*. 1995; Chu and Herskowitz 1998).

In this study we have identified Nup2 in an enriched protein pool that co-purifies with tagged Ndj1 in a yeast meiotic extract. We show that Nup2 is required for normal spore viability and meiotic progression and we map this function to a 125 amino-acid region of the 720 amino acid protein. This region we call the <u>m</u>eiotic <u>a</u>utonomous <u>r</u>egion (MAR) is both necessary and sufficient for Nup2’s meiotic function and the MAR alone is enriched at the nuclear periphery. A number of synthetic *nup2Δ ndj1Δ* defects suggest Nup2 and Ndj1operate in different pathways. We propose that a chromosome organizing function of Nup2 acts in concert with the Ndj1-mediated bouquet to promote the organizational features of the nucleus that can support the dynamic chromosome events of meiotic prophase.

## MATERIALS and METHODS

### Strains

All strains in this study are derivatives of SK1 and listed in Table 2. All media were generated as previously described (Ho and Burgess 2011; Lui *et al*. 2013). Gene knockouts and fluorescent tagging were constructed using standard tailed PCR based gene replacement and tagging techniques (Longtine *et al*. 1998; Goldstein and McCusker 1999; Sheff and Thorn 2004; Lee *et al*. 2013). All gene knockouts were confirmed by PCR and new alleles were confirmed by DNA sequencing. Double and triple mutants were created by tetrad dissection. All GFP and mRuby2 protein tags were added to the C-terminal end of the tagged protein with the exception of Zip1. Zip1 was tagged with GFP internally by integrating a ZIP1-GFP plasmid marked with *URA3* at the endogenous *ZIP1* locus and looping out the endogenous *ZIP1* (Scherthan *et al*. 2007). Truncations were generated using 2-step allele replacement (Rothstein 1991). Genomic preparations for PCR were generated as previously described (Danilevich and Grishin 2002).

### Meiotic time course protocol

The time course protocol was followed as previously reported (Ho and Burgess 2011; Lui *et al*. 2013). Cells were synchronized for progression through meiosis by first patching and mating haploid cells taken from glycerol stocks stored at – 80^o^ C on YP media plates (2% Bacto peptone, 1% Bacto yeast extract, 0.01% adenine sulphate, 0.004% tryptophan, 0.002% uracil) supplemented with 3% glycerol and 2% bacto agar, for 15 hrs at 30°C. Cells were then streaked onto YPD plates (YP plus 2% dextrose, 2% bacto agar) for 2 days at 30° C. Well-isolated diploid single colonies were used to inoculate 5ml liquid YPD cultures (YPD w/o uracil supplementation) and incubated for 30 hrs at 30° C on a roller drum. The YPD liquid culture was then added to YPA (YP + 1% KOAc w/o uracil supplementation) to a final OD600 of 0.23 and grown for 14.5 hrs at 30° C on a roller drum. Cells were pelleted by brief centrifugation, washed with SPM (1% KOAc, 0.02% raffinose, 0.009% SC dropout powder) and resuspended in SPM to a final OD600 of 3.0. Cells were removed from the culture at various time points thereafter (starting at t = 0 hrs), fixed in 40% ethanol and stained with DAPI to follow the staged kinetics of meiotic prophase (e.g. synapsis, DSB formation and repair, and nuclear divisions. Nuclear division was marked by the formation of cells with two well-differentiated DAPI stained foci. At least 200 cells were analyzed for each time point. Lifespan analysis was carried out as previously described (Padmore *et al*. 1991; Wanat *et al*. 2008).

### Purification of Ndj1-TAP tagged proteins

Ndj1 was C-terminally TAP tagged as previously described (Rigaut *et al*. 1999). Ndj1 was purified as previously described with modifications (Puig *et al*. 2001). Four liters of meiotic time course culture were harvested after four hrs when bouquet formation peaks (Wanat *et al*. 2008). The cell pellet was resuspended in lysis buffer (50mM HEPES pH 7.5, 150mM NaCl, 20% glycerol, 0.1% NP-40, 1mM DTT, 1mM PMSF, 10mM sodium pyrophosphate, 50mM NaF, 60mM β-glycerophosphate, 1X Halt Protease Inhibitor Cocktail (ThermoFisher Scientific; Conrad *et al*. 2008). The resuspended cells were bead beat using a Beadbeater Disrupter (Biospec) with 0.5 mm diameter zirconia/silica beads (Biospec) with eight cycles of one minute of bead beating followed by two minutes of on ice. The cell homogenate was spun at 45000 × g for 30 minutes at 4° C. The supernatant was removed and mixed with 0.5ml of IgG sepharose 6 fast flow beads (GE) preequilibrated with lysis buffer for three hrs at 4° C. The bead slurry was then run through a 0.8 × 4-cm Poly-Prep column (BioRad). The beads were then washed three times with 10ml of lysis buffer and once with 10ml of TEV cleavage buffer (10mM Tris pH8.0, 150mM NaCl, 10% glycerol, 0.5mM EDTA, 0.1% NP-40, 1mM DTT). The beads were then digested in 1ml of TEV cleavage buffer with 100 units of AcTEV protease (ThermoFisher Scientific) rotating overnight at 4° C. The digested products were then eluted and mixed with 3ml of calmodulin binding buffer (10mM Tris pH8.0, 150mM NaCl, 1mM imidazole, 1mM MgOAc, 2mM CaCl, 10% glycerol, 10mM β-mercaptoethanol, 0.1% NP-40) and 3µl of 1M CaCl. The solution was then added to 0.5ml of Calmodulin Affinity Resin (Agilent) preequilibrated with calmodulin binding buffer in a 0.8 × 4-cm Poly-Prep column (BioRad) and mixed for two hrs at 4° C. The beads were then washed three times with 10ml of calmodulin binding buffer. Following the wash the beads were directly digested and analyzed by the UC Davis Proteomics core for mass spectroscopy. Two experimental samples with Ndj1 TAP tagged and one negative wild-type control without Ndj1 tagged were analyzed the results are reported in File S1.

### Spore viability and nondisjunction analysis

Spore viability was determined as the percent viable spores from dissected tetrads. Single colonies from YPD plates were patched onto solid SPM media and incubated for three days at 30° C, followed by spore dissection on YPD plates. For each strain, spore viabilities are from pooled tetrad data dissected from multiple colonies on multiple independent occasions. Sporulation efficiency was scored as a cell having two or more spores. A computational estimation of nondisjunction frequencies was determined using TetFit (Chu and Burgess 2016). Expected live:dead tetrad frequencies were determined using the R-suite (TetSim, TetFit, and TetFit.Test) with the following conditions: For TetFit, the number of nondisjunction intervals (ndint) was 3000, the number of random spore death intervals (rsdint) was 3000, ANID was 0.035, and the MI-ND multiplier (ndm) was 10. Detailed background and instructions and examples of the outputs for using the R-scripts are provided in File S2. For multiple comparisons the p values were corrected using the Benjamini and Hochberg method (Benjamini and Hochberg 1995).

### Spotting assay for vegetative growth

Diploid cells were prepared for synchronization as described above except that cells from the five ml YPD culture were diluted to an OD600 = 1.0 after 24 hrs growth. Suspended cells were serially diluted 1:10 in a 96-well plate and 2.5 µl of the diluted cultures were spotted onto YPD and minimal medium plates using a multichannel pipettor, followed by incubation at 20° C, 25° C, 30° C, and 37° C. YPD plates were imaged after incubation for 48 and 72 hrs and minimal medium plates (0.67% Difco yeast nitrogen base, 2% dextrose, 2% agar) were imaged after incubation for 72 and 120 hrs using an Epson Perfection 1200 U scanner.

### Cell fixation and imaging

A 0.5ml aliquot of cultured cells were pelleted by centrifugation and resuspended in one ml of ice-cold fixative (1% formaldehyde, 100mM KPO4 pH 7.5, 4% sucrose) followed by incubation at 4° C for 30 min while gently rotating. Cells were washed twice with one ml of ice-cold KPO4/sorbitol solution (1.2 M sorbitol, 100mM KPO4 pH 7.5, 0.01% Na-Azide) and resuspended in 100 µl KPO4/sorbitol solution. Slide preparation for imaging cells was performed as previously described (Dresser 2009). Cells were imaged between an SPM agar pad and cover slip. Imaging of TetR-GFP bound at the *URA3::tetOx224* locus for the pairing assay was performed as previously described (Lui *et al*. 2013).

A Hybrid Marianas confocal spinning disk 3D fluorescence wide-field microscope (Intelligent Imaging Innovations, 3i) was used for imaging using a 100 × 1.46 NA oil objective lens (Olympus) and with a Yokogawa spinning disk head at room temperature. Fluorescence microscopy data was acquired using an electron multiplying charge coupled device camera. All images were analyzed using ImageJ and SlideBook (3i software) and are shown as a single Z-slice or projections of no fewer than 23, 233 nm Z-steps. The fluorophores imaged in this paper were DAPI, eGFP, and mRuby2 (Sheff and Thorn 2004; Lee *et al*. 2013). At least 200 cells were analyzed per time point for all assays assessing proportion of cells progressing through meiosis.

### DNA analysis

DNA physical assays and meiotic progression analysis were performed as described previously (Oh *et al*. 2009). Analysis of CO and NCO products was carried out by digesting purified genomic DNA with XhoI or XhoI and NgoMIV, respectively (Oh *et al*. 2009; Wu *et al*. 2010). The reported CO and NCO are based on measurements from three independent colonies. The mean and standard deviation are reported and p values were corrected for mutliple comparisons using the “Holm” method in R (Holm 1979).

### Data availability

Strains are available upon request. File S1 is an Excel spreadsheet showing the enrichment of proteins analyzed by mass spectrometry. File S2 contains the R-scripts and detailed instructions for using TetSim, TetFit, and TetFit.Test from Chu and Burgess, 2016).

## RESULTS

### Identification of a role for Nup2 in normal meiotic progression

We identified five candidate proteins that co-purified with a C-terminal TAP-tagged Ndj1 protein by mass spectrometry from mid-prophase meiotic cell extracts: Rif1, Esc8, Yrf1, Nop56, and Nup2 (Figure 1, B-C; File S1). All five proteins were shown previously to localize in the nucleus in vegetatively growing cells (Huh *et al*. 2003). Rif1 is a nonessential protein that binds to the C-terminal region of Rap1 and is involved in telomere silencing and regulation of telomere length (Wotton and Shore 1997; Teixeira *et al*. 2004). Esc8 is a nonessential protein involved in telomeric and mating-type locus silencing and interacts with Sir2 (Cuperus and Shore 2002). *YRF1* is a repeated gene encoded by the Y’ element of subtelomeric regions, is highly expressed in mutants lacking the telomerase component *TLC1, and* the protein product is potentially phosphorylated by Cdc28 (Yamada *et al*. 1998; Ubersax *et al*. 2003). We did not characterize a possible role of Yrf1 since it has multiple paralogs. Nop56 is an evolutionarily-conserved component of the box C/D snoRNP complex (Lafontaine and Tollervey 2000). Since Nop56 is essential, we did not explore a possible role in meiosis. Nup2 is a component of the nuclear pore complex (NPC), exhibits boundary activity, and binds at hundreds of sites throughout the genome (Loeb *et al*. 1993; Dilworth *et al*. 2005; Schmid *et al*. 2006; Light *et al*. 2010). None of these proteins has been previously shown to interact physically with Ndj1.

We tested if deleting *RIF1, ESC8,* or *NUP2* decreased sporulation efficiency, spore viability, or delayed meiotic progression, which are observed in *ndj1Δ* (Table 1). In all cases, the mutants sporulated with near wild-type (WT) efficiency (> 91%, n = 200), yet only the *nup2Δ* mutation conferred decreased levels of spore viability compared to WT (86.1%, n = 1,536 and 97.1%, n = 1152, respectively; Table 1). To measure the kinetics of meiosis I division (MI), we calculated the fraction of multinucleate cells taken from a synchronized cell culture for various hours following transfer to sporulation medium starting at t = 0 hrs. The *rif1Δ and esc8Δ* mutants progressed through MI at the same rate as WT (Figure 1D). Since *RIF1 and RIF2* are partially redundant (Wotton and Shore 1997), we analyzed r*if2Δ and* the *rif1Δ rif2Δ* double mutant and also found no delay in nuclear divisions (Figure S1). Interestingly, the formation of multinucleated cells in the *nup2Δ* mutant was delayed by ∼ 2 hrs compared to WT, which is similar to the *ndj1Δ* phenotype (Figure 1D; Conrad *et al*. 1997). We performed a colony growth assay to test if *nup2Δ* gave decreased grow efficiency, which might cause asynchrony in the cultures and appear to delay MI progression. A spotting assay of serially diluted cells showed that *nup2Δ* growth was no less efficient than WT on either rich or minimal medium at low or high temperatures (Figure 1G). Growth efficiency of *nup2Δ* on YPD at 37° C was slightly more efficient than WT. These data point to a role for Nup2 in promoting normal meiotic progression that is dispensable for normal mitotic growth.

**Table 1.** Sporulation efficiency and spore viability for strains used in this study

| Strain | % Sporulation efficiency (n) | Tetrads dissected | 4-sv | 3-sv | 2-sv | 1-sv | 0-sv | % Spore viability |
| --- | --- | --- | --- | --- | --- | --- | --- | --- |
| WT | 95.0% (200) | 288 | 269 | 10 | 6 | 1 | 2 | 97.1 |
| WT<sup>a</sup> | 89% (231) | 1199 | 1087 | 77 | 32 | 1 | 2 | 96.8 |
| *nup2Δ*<sup>b</sup> | 92.0% (200) | 384 | 239 | 95 | 36 | 10 | 4 | 86.1 |
| *esc8Δ* | 91.5% (200) | 48 | 42 | 3 | 2 | 1 | 0 | 94.8 |
| *rif1Δ* | 94.0 (200) | 48 | 41 | 4 | 2 | 0 | 1 | 96.7 |
| *nup53Δ* | 94.0% (200) | 72 | 66 | 3 | 1 | 1 | 1 | 95.8 |
| *nup100Δ* | 93.0% (200) | 72 | 69 | 2 | 1 | 0 | 0 | 95.9 |
| *nup157Δ* | 94.5% (200) | 72 | 69 | 1 | 2 | 0 | 0 | 98.2 |
| *nup2Δ557-720* | 92.5% (200) | 48 | 42 | 4 | 2 | 0 | 0 | 95.8 |
| *nup2Δ176-720* | 91.5% (200) | 48 | 45 | 2 | 1 | 0 | 0 | 97.9 |
| *nup2Δ2-50* | 93.0% (200) | 72 | 67 | 2 | 3 | 0 | 0 | 97.2 |
| *nup2Δ2-175* | 94.0% (200) | 48 | 28 | 12 | 8 | 0 | 0 | 84.5 |
| *nup2-MAR* | 92.5% (200) | 96 | 83 | 11 | 2 | 0 | 0 | 96.1 |
| *nup2-MAR-GFP* | 94.0% (200) | 48 | 45 | 2 | 1 | 0 | 0 | 97.9 |
| *nup60Δ* | 71% (200) | 264 | 85 | 96 | 46 | 25 | 12 | 70.6 |
| *htz1Δ* | 67.0% (200) | 288 | 77 | 91 | 69 | 40 | 11 | 66.0 |
| *ndj1Δ* | 75.0% (200) | 192 | 136 | 14 | 19 | 6 | 17 | 82.0 |
| *ndj1Δ*<sup>a</sup> | 59% (221) | 851 | 482 | 112 | 123 | 38 | 96 | 74.9 |
| *nup2Δ ndj1Δ* | 1.5% (200) | 144 | 83 | 40 | 15 | 6 | 0 | 84.7 |
| *nup53Δ ndj1Δ* | 75.0% (200) | 144 | 103 | 10 | 12 | 6 | 13 | 82.0 |
| *nup100Δ ndj1Δ* | 74.5% (200) | 72 | 48 | 3 | 14 | 1 | 6 | 79.9 |
| *nup157Δ ndj1Δ* | 72.5% (200) | 72 | 39 | 5 | 14 | 4 | 10 | 70.5 |
| *csm4Δ* | 77.5% (200) |  |  |  |  |  |  |  |
| *nup2Δ csm4* | 1.0% (200) |  |  |  |  |  |  |  |
| *spo11Δ* | 93.5% (200) |  |  |  |  |  |  |  |
| *nup2Δ spo11Δ* | 0.5% (200) |  |  |  |  |  |  |  |

<sup>a</sup> Data from Wanat et al., 2008 for comparison.
<sup>b</sup> Tetrad data from SBY3945 (n = 98) and SBY4102 (n = 288) were pooled.

**Table 2.**
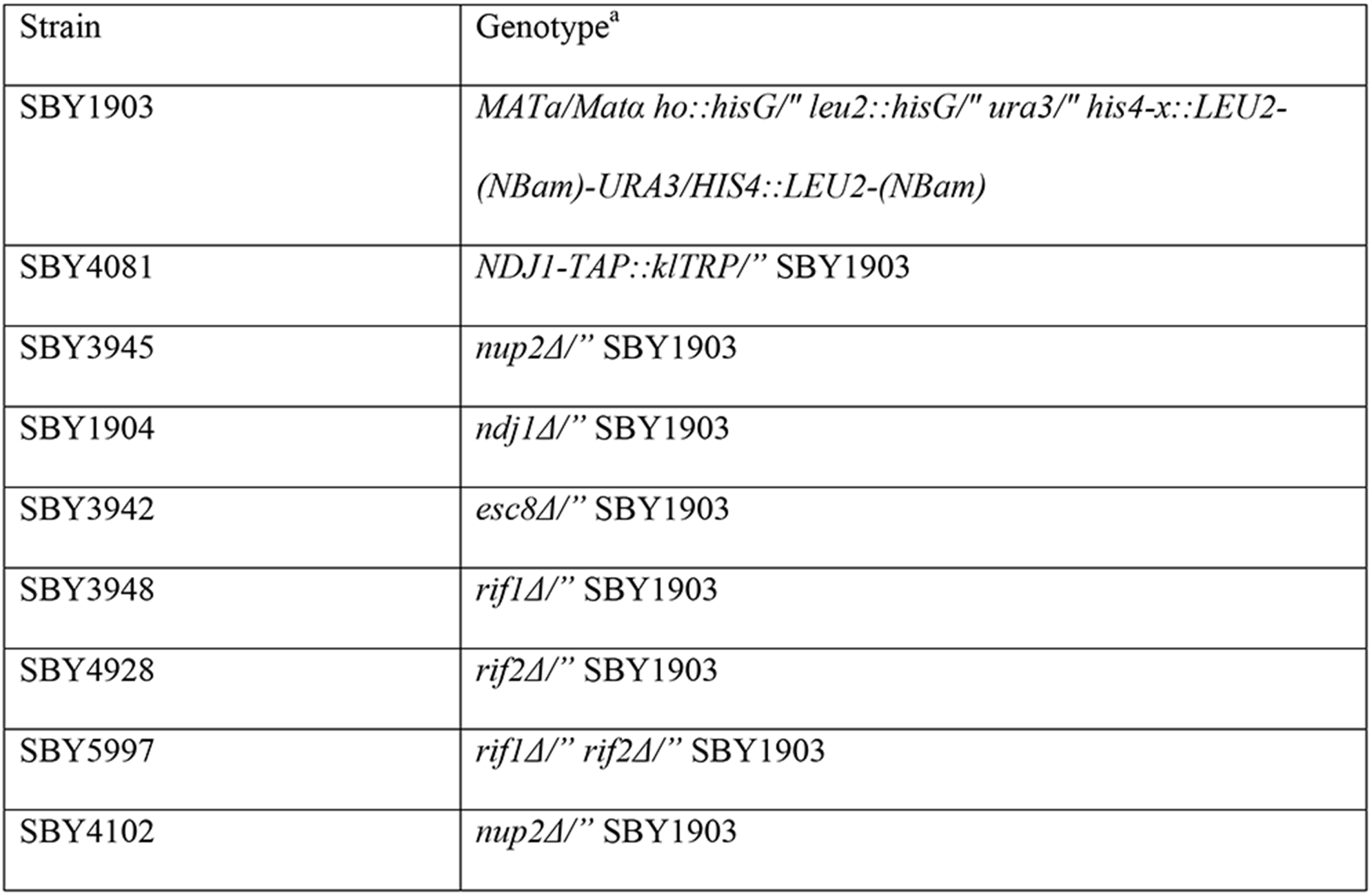
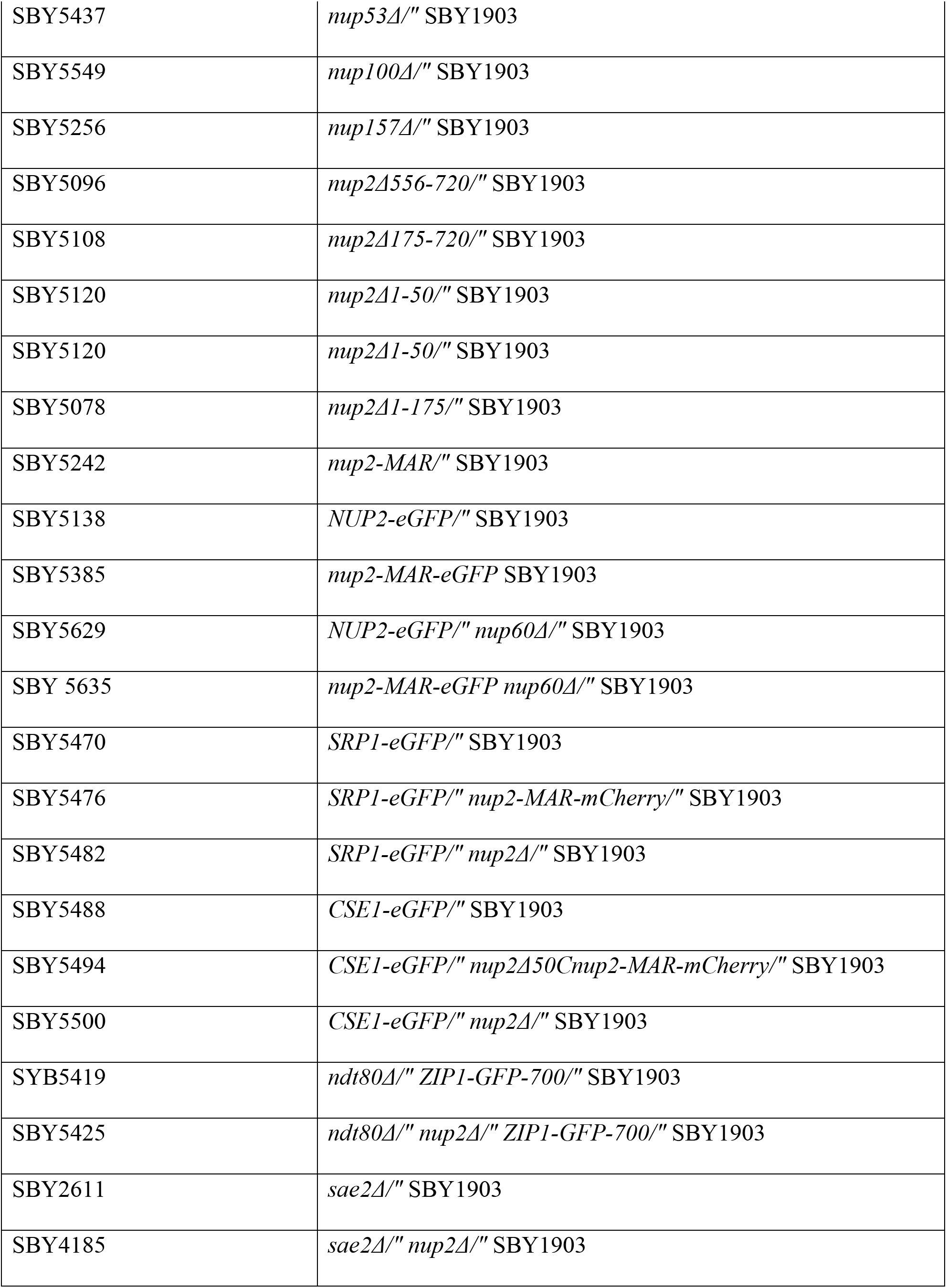
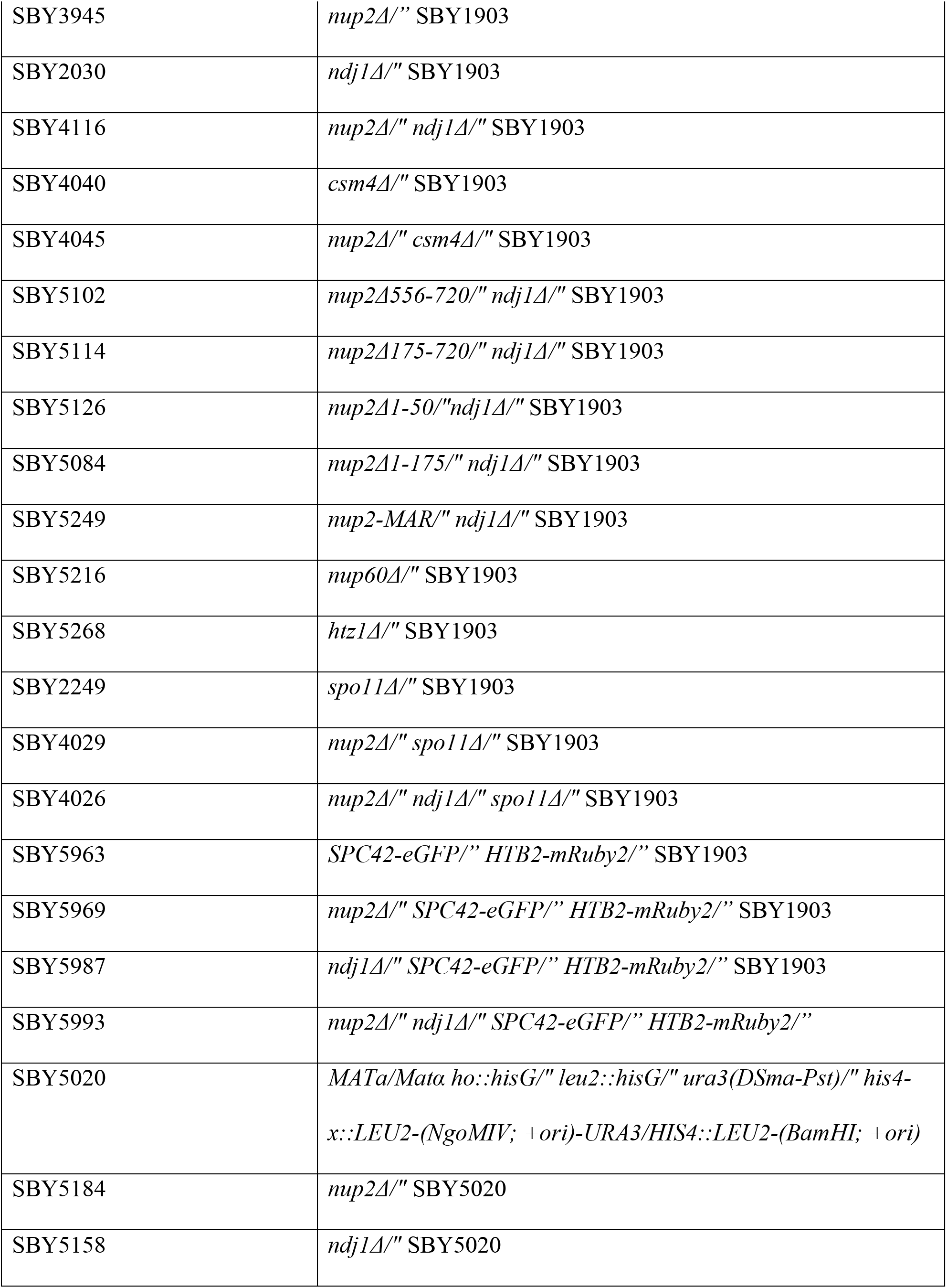
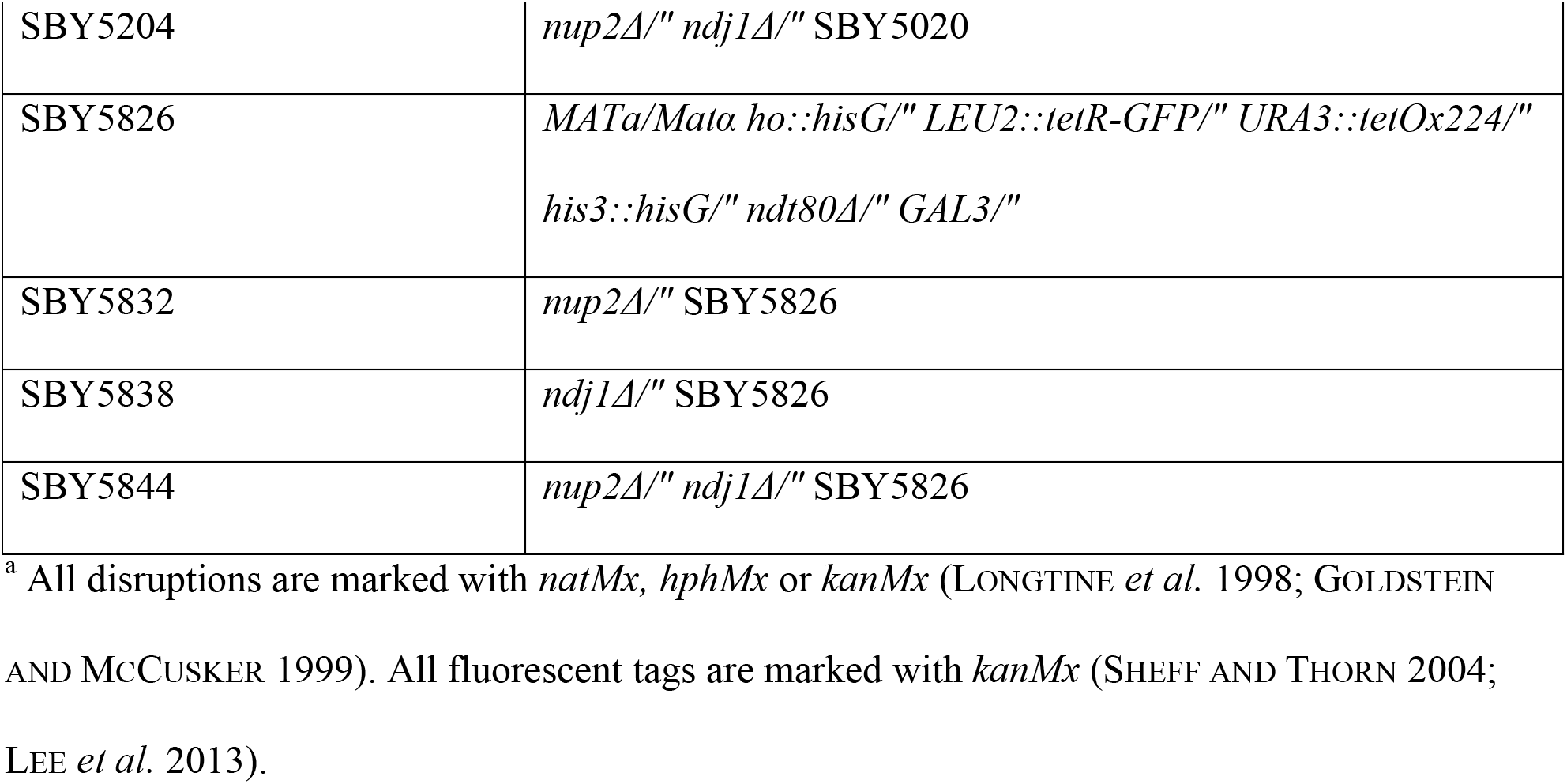
Yeast strains used in this study

### Meiotic phenotypes for other *nup* mutants

Other *nup* deletions (*nup53Δ, nup100Δ and nup157Δ*) showed no decrease in spore viability, delayed meiotic progression, or other obvious growth defects (Figure 1, E and F; Figure S2). The *nup60Δ and nup84Δ* mutants showed more severe growth defects at 30° C and were excluded from time-course analysis due to the low likelihood that these cells could be synchronized in culture. While 70% of *nup84Δ* cells could undergo at least one nuclear division, less than 1% of cells gave two or more spores (n = 200 for both). Sporulation efficiency and spore viability were both reduced in *nup60Δ*, and like *nup2Δ,* it showed more efficient growth than WT at 37° C (Table 1 and Figure S2). Together, these data suggest that the nonessential nucleoporins Nup2, Nup60, and Nup84 are required for normal meiosis. Given its relationship to Ndj1 and Nup60 (below), we focused our attention on Nup2.

### A 125 amino-acid domain of Nup2 is both necessary and sufficient for its role in promoting meiotic progression and spore viability

To test if the *nup2Δ* meiotic phenotypes are due to a nuclear transport defect(s), we created a series of truncation mutations in one or more of Nup2’s three characterized transport domains (Figure 2A): 1) the first 50 amino acid N-terminal domain, which binds Srp1/Kap60 (importin-α; Hood *et al*. 2000; Matsuura *et al*. 2003), 2) the 154 amino acid C-terminus (aa 557-720) that binds the Ran-GTP homolog, Gsp2 (Booth *et al*. 1999; Hood *et al*. 2000; Matsuura *et al*. 2003), and 3) the 381 amino acid unstructured domain containing FXFG repeats, which binds to Kap95 (importin-β; aa 176-556; Solsbacher *et al*. 2000). We found that all three functional domains of Nup2 could be deleted either on their own or together without negatively affecting spore viability (Figure 2A). Deleting the region encoding the first 175 amino acids of Nup2 (*nup2Δ2-175*) was the only other deletion mutation that conferred a *nup2Δ*-like meiotic phenotype. Together, the deletion mapping pointed to an uncharacterized 125-amino acid region of the protein likely to have a role in meiosis. We expressed this region alone and found it was both necessary and sufficient for supporting Nup2’s role in promoting spore viability and meiotic progression (Figure 2A, below). We refer to this region as the Nup2-<u>m</u>eiotic <u>a</u>utonomous <u>r</u>egion (Nup2-MAR).

**Figure 2.**
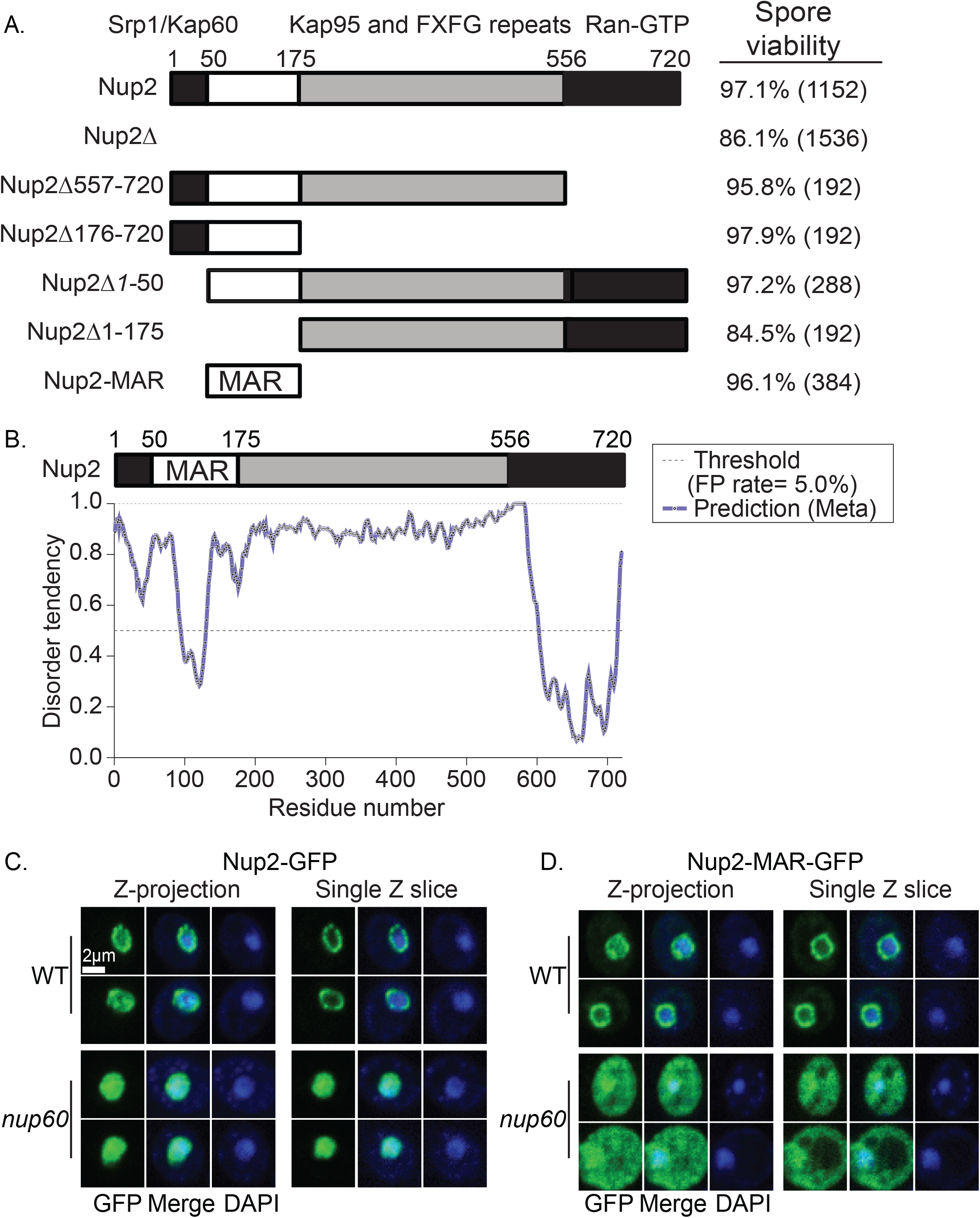
Mapping the functional meiotic-autonomous region of Nup2 and its localization in fixed cells. **A**. chematic of the 720 amino acid Nup2 protein architecture and deletion of known functional domains involved in nucleocytoplasmic transport. The known protein-binding partners that have been associated with specific regions of Nup2 are indicated. All strains are in the SBY1903 background; *nup2Δ557-720* (SBY5096), *nup2Δ176-720* (SBY5108), *nup2Δ1-50* (SBY5120), *nup2Δ1-175* (SBY5078), *nup2-MAR* (SBY5242).The spore viability and the number of spores analyzed from four-spore tetrads are given on the right. **B**. Disorder profile plots using metaPrDOS of the full length Nup2 protein. A higher score indicates a region more likely to be disordered. The dashed line represents the 5% cutoff for false positives. **C**. Localization of Nup2-GFP in fixed cells taken from a mitotically dividing culture. Fixed cells were stained with DAPI and imaged using confocal spinning disk 3D fluorescence wide-field microscope. Two representative cells are shown for WT (SBY5138) and *nup60Δ* (SBY5629) strains. Left: Images are maximum-intensity projections from no fewer than 23, 233 nm Z-steps. Right: Images are from a single Z-slice taken from about the middle of the nucleus. **D**. Localization of the 125-amino acid Nup2-MAR-GFP region as described in part C in WT (SBY5385) and the *nup60Δ* (SBY5635).

### Nup2-MAR likely folds into a discrete protein domain

A protein BLAST search of the Nup2-MAR sequence returns Nup2 orthologs from related fungal species and weak matches to the Nup2 mammalian ortholog Nup50. No domains or motifs were found using the PROSITE or Pfam databases of protein families and domains (Finn *et al*. 2010; Sigrist *et al*. 2010). The MAR sequence is predicted to form a stable structure based on the Meta Protein DisOrder prediction System (metaPrDOS; Figure 2B; Ishida and Kinoshita 2008).

### Nup2-MAR enrichment at the nuclear periphery depends on Nup60

Nup2 is a mobile nucleoporin that is enriched near the inner-nuclear envelope via binding to Nup60 (Hood *et al*. 2000; Solsbacher *et al*. 2000), but is also present in the nucleoplasm (Loeb *et al*. 1993; Denning *et al*. 2001). To test if the Nup2-MAR exhibits a similar nuclear location pattern as Nup2, we expressed Nup2-GFP and Nup2-MAR-GFP and analyzed their intracellular localization in fixed cells from an exponentially dividing culture. We found that both Nup2-GFP and Nup2-MAR-GFP signal was enriched at the nuclear periphery (Figure 2C). The Nup2-MAR-GFP fusion protein also showed some diffuse signal in the cytoplasm not seen for Nup2-GFP, which would be expected due to its small size allowing it to diffuse across the NPC (41 kDa; Figure 2C; Shulga *et al*. 2000). Previous studies have demonstrated that Nup2’s localization to the nuclear basket requires Nup60; however, there are also low levels of Nup2 free of the nuclear basket in the nucleoplasm (Denning *et al*. 2001). We tested the cellular localization of Nup2-GFP and Nup2-MAR-GFP in a *nup60Δ* mutant background and found in both cases the nuclear signal was diffuse, with no obvious enrichment at the nuclear periphery. Together, these data suggest that the Nup2-MAR sequence is sufficient to target Nup2 to the nuclear periphery, and that this localization requires Nup60.

Nup60 and Htz1 have been shown previously to act with Nup2 in promoting boundary activity at subtelomeric regions near the nuclear envelope (Dilworth *et al*. 2005). We thus tested if deletion of *NUP60* or *HTZ1* gives a similar meiotic defect as *nup2Δ* in meiosis. We found that both exhibited reduced sporulation efficiency (71%, n = 200 and 67%, n = 200) compared to *nup2Δ* (92%, n = 200) as well as reduced spore viability (70.6%, n = 1056 and 66.0%, n = 1152) compared to *nup2Δ* (86.1%; Table 1). Thus, while Nup60 is required to localize Nup2 to the nuclear periphery, it may also disrupt meiosis in independent ways.

### The MAR sequence is required for normal meiotic progression independent of transport function

Since the Nup2-MAR is missing every known sequence element associated with transport function, we expected that Nup2’s contribution to transport function would be disrupted in the *nup2-MAR* mutant. In a previous study, both *nup2Δ50* (an N-terminal deletion missing the Srp1/Kap60 binding domain and the MAR) and *nup2Δ* mutants were found to be defective for the enriched localization of Srp1 and Cse1 near the nuclear envelope (Booth *et al*. 1999; Solsbacher *et al*. 2000; Matsuura *et al*. 2003). We expected this would also be the case for the *nup2-MAR* allele. By tagging Srp1/Kap60 and Cse1 with GFP, we found that these proteins localized to the nuclear periphery, with some cytoplasmic and nuclear signal in WT cells. This localization was disrupted in both the *nup2Δ and nup2-MAR* mutants since both protein fusions shifted to a more diffuse staining pattern in the nucleus (Figure S3). These results support the notion that Nup2 (and Nup2-MAR) function(s) during meiosis is independent of Nup2’s role in nucleocytoplasmic transport.

### The onset of zygotene, but not its duration, is delayed in *nup2Δ*

We next examined if meiotic prophase events preceding nuclear division are delayed in the absence of Nup2. Synapsis between homologous chromosomes in budding yeast is initiated by the formation of DSBs which are required for the formation of crossovers that ensure homolog disjunction at MI (Zickler and Kleckner 2015). Synapsis initiates during zygotene while DSBs are undergoing repair and is complete by pachytene. To monitor entry into zygotene, we examined SC formation through analysis of a Zip1-GFP fusion protein (Scherthan *et al*. 2007). Cells in zygotene exhibit “patchy” GFP signal as they transition to full synapsis at pachytene when the Zip1-GFP signal appears as lines (Figure 3, A and B). To determine the fraction of cells that reach pachytene, we deleted *NDT80*, a regulatory gene required for cells to progress beyond the pachytene stage (Xu *et al*. 1995).Using a meiotic time-course assay as described above, we found that the peak occurrence of zygotene in WT and *nup2Δ* was marked at three and four hrs, respectively (Figure 3A). By 12 hrs, 99% of *ndt80Δ and* 94% of *nup2Δ ndt80Δ* cells reached the pachytene stage.

**Figure 3.**
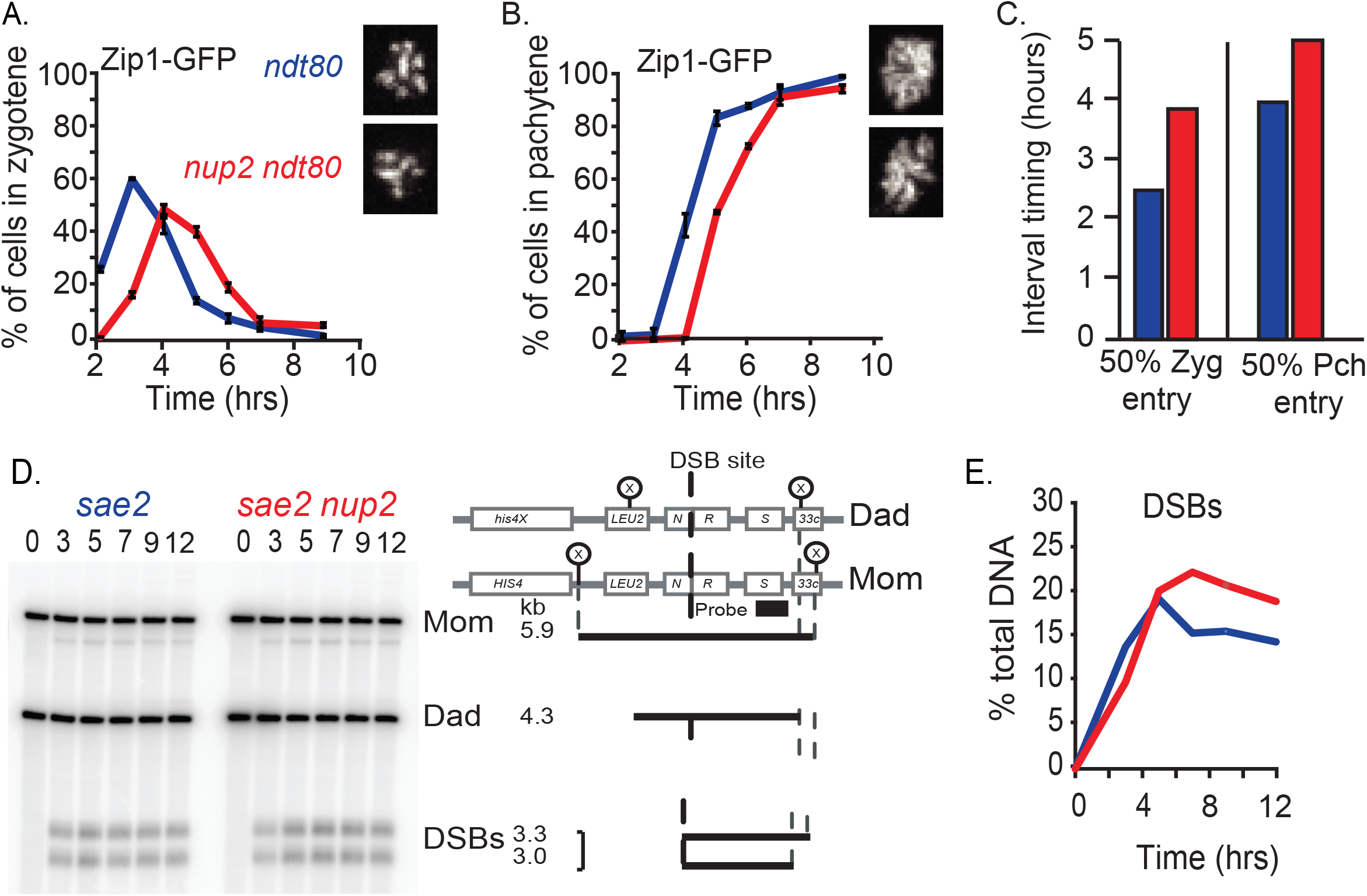
Delay of entry into zygotene and pachytene and increased levels of DSBs in the absence of Nup2. **A-B**. Meiotic time course of *ntd80Δ* (Blue; SYB5419) and *nup2Δndt80* (Red; SBY5425) strains expressing Zip1-GFP. Shown is the fraction of cells (n = 200) scored as Zygotene (A) and pachytene (B). Representative maximum intensity Z-projections of image stacks are shown on the right side of each figure. Error bars represent the average ± SD for three cultures run in parallel. Similar results were seen for time courses run on different days. **C**. The calculated zygotene entry and lifespans from (A and B) of *ntd80Δ* (blue), *nup2Δndt80* (red). The life spans of these events (the time between which 50% of cells enter and 50% of cells exit) each phase were calculated. **D**. outhern blot of DNA isolated from *sae2Δ* (SBY2611) and *nup2Δ sae2Δ* (SBY4185) mutant cells in a meiotic time courses. Left: Migration of DNA products of a XhoI digest of genomic DNA after gel electrophoresis and probed with sequences that reside on the right side of the DSB hot spot after transfer to a nylon membrane. **E**. Quantifications of DSB products (DSBs/Total DNA) from the Southern blot depicted in part (D) of *sae2Δ* (blue) and *nup2Δ sae2Δ* (red). Both DSB bands were summed to calculate DSB as a percent of total DNA in the lane (Right). Data from one experiment is shown. Identical results were found in an independent replica experiment done on a different day.

Based on these data, we next calculated the time window, or lifespan, in which WT and mutant cells transit through zygotene to pachytene (Padmore *et al*. 1991). The lifespan of the zygotene stage can be derived from the area under the curve of the primary data (Hunter and Kleckner 2001; Wanat *et al*. 2008). An increase in lifespan compared to WT indicates a specific delay in passing through the corresponding stage (Hunter and Kleckner 2001). The calculated lifespans for zygotene in *ndt80Δ and nup2Δ ndt80Δ* show that deleting *NUP2* does not cause a delay in this stage (1.7 and 1.5 hrs, respectively; Figure 4C). Therefore, while zygotene entry was delayed by at least one an hour in the absence of Nup2, the time cells took to transit from zygotene to pachytene was not delayed.

**Figure 4.**
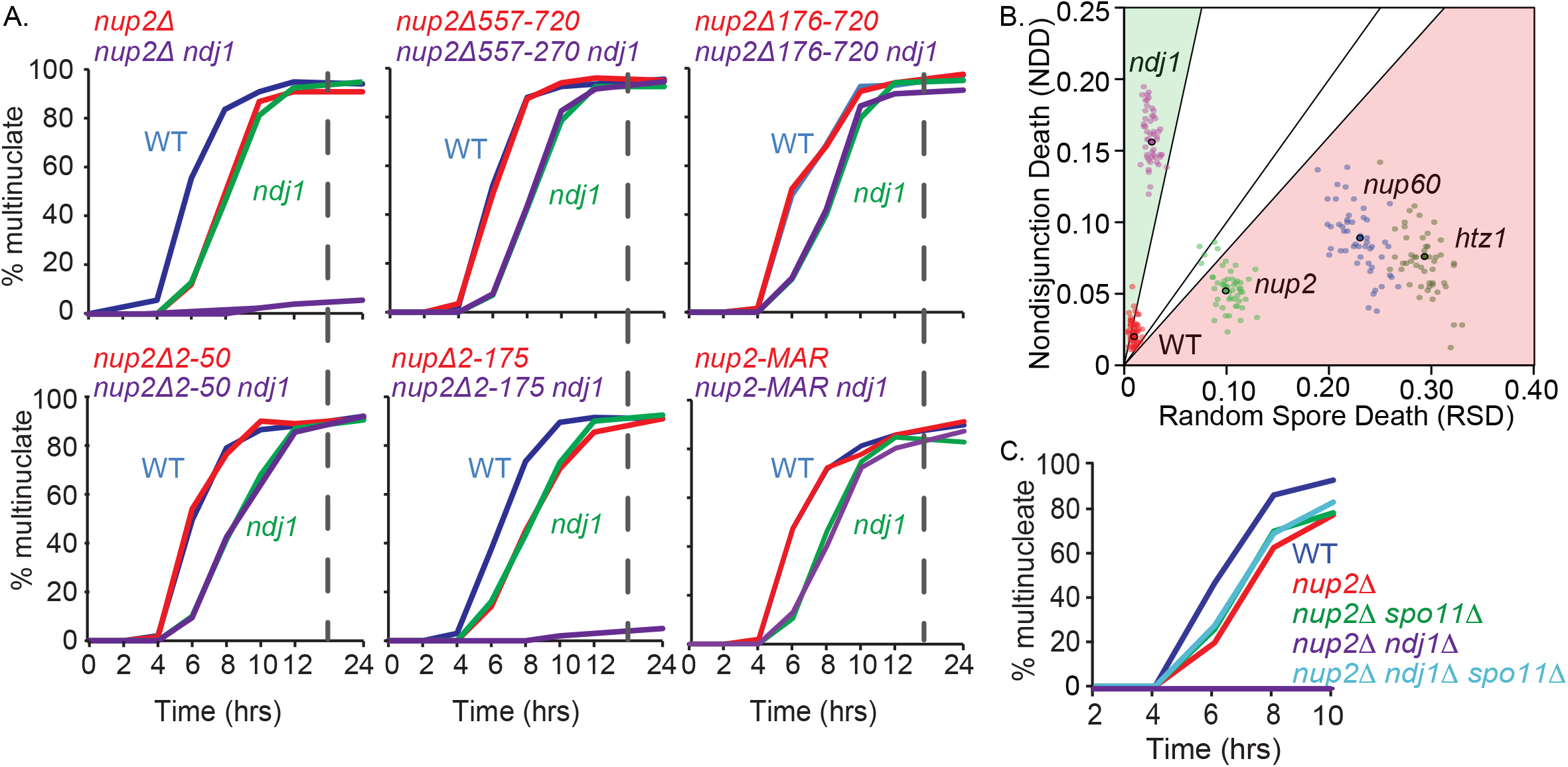
Synthetic block to nuclear division in mutants carrying both a *nup2Δ* truncation mutations and *ndj1Δ*. **A**. All truncation mutations where the MAR is present progress past meiosis I with no delay compared to WT in meiotic time course experiments described in Figure 1D. Allele names are the same as shown in Figure 2. Only *nup2Δ* (SBY4102) and *nup2Δ2-175* show a dramatic reduction in the percent of multinucleate cells even after 24 hrs in SPM. This phenotype is complemented by a truncation *nup2-MAR* in which all amino acids of Nup2 except for 51-175 are deleted. **B**. ause of spore death in WT (SBY1903) *ndj1Δ* (SBY2030), *nup2Δ* (SBY4102 and SBY3945), *nup60Δ* (SBY5216) and *htz1Δ* (SBY5268). TetFit was used to calculate the estimated contributions of spore death due to nondisjunction (NDD) and by random spore death (RSD), which gives the best fit to the experimental data set (Chu and Burgess, 2016). In this assay precocious sister-chromatid separation or defects in MII division would appear as RSD. A detailed description of the TetSim and TetFit R-Scripts with annotated instruction is in Sup File 2. To analyze the variation of the output values from TetFit-A, we ran 50 independent simulations using TetFit.Test using the calculated NDD and RSD values for each genotype and the corresponding number of experimentally dissected tetrads (Figure 4B, E columns; Table S2). The graphical output of the simulations in Figure 4C shows that each genotype gives well defined clusters representing NDD and RSD values for 50 simulated tetrad data sets. **C**. Kinetics of nuclear division in a time course experiment with WT, *nup2Δ* (SBY3945), *nup2Δ spo11Δ* (SBY4029), *nup2Δ ndj1Δ* (SBY3983), and *nup2Δ ndj1Δ spo11Δ* (SBY4026).

### DSBs formation in the absence of Nup2

Since synapsis initiates at the sites of DSBs, we tested if the delay entering the zygotene stage corresponded to a reduction or a delay in the formation of DSBs. To test this we measured the physical precursors and products of DSB formation at the well-characterized *HIS4LEU2* hot-spot locus (Cao *et al*. 1990; Storlazzi *et al*. 1995; Hunter and Kleckner 2001). Following restriction digestion of DNA isolated from cells in a meiotic time course, the DNA fragments were resolved and probed by Southern blotting.

The predicted migration of bands reflects the precursor, intermediates and products of the formation and repair of Spo11-induced DSBs. For this experiment, we analyzed *NUP2 sae2Δ and nup2Δ sae2Δ* cells where the resection of DSBs is prevented. This gives a readout of total DSBs formed without being turned over (McKee and Kleckner 1997).We found that peak DSB levels occurred at five and seven hrs for *sae2Δ and nup2Δ sae2Δ* strains, respectively. In the *sae2Δ* mutant, DSB levels reached ∼20% of total DNA levels, while in the *nup2Δ sae2Δ* mutant, peak levels were 23% of total DNA (Figure 4, D and E). At later time points, the *nup2Δ sae2Δ* mutant also gave overall greater levels of DSBs compared to *sae2Δ*. A second time course experiment gave a similar result. The cause of the early delay phenotype was not explored further, yet it is consistent with the possibility that the *nup2Δ* mutation may disrupt meiotic DNA replication or early chromosome axis structure (Blitzblau and Hochwagen 2013).

### The *nup2Δ ndj1Δ* mutant exhibits a synthetic sporulation defect and a failure to form binucleate cells

Since we identified Nup2 as a possible interactor of Ndj1, we tested if Nup2 and Ndj1 act in the same or different pathways to promote timely meiotic progression.

Interestingly, while 92% of *nup2Δ and* 75% of *ndj1Δ* cells gave two or more spores, only 5% of *nup2Δ ndj1Δ* double mutant cells sporulated (Table 1). The synthetic sporulation phenotype was also seen for *nup2Δ1-175* mutant, also lacking the MAR domain, but not in the other deletion strains where the MAR was present, including *nup2-MAR* (Table 1).

Both the *nup2Δ and ndj1Δ* mutants gave a modest reduction in spore viability among 4-spore tetrads (86.1%, n = 1,536, and 82.0%, n = 768, respectively) compared to WT (97.1%, n = 1152). Of the 1% spores formed from the *nup2Δ ndj1Δ* double mutant, the spore viability was 84.7% (n = 576). While these values could be highly skewed from the very low sporulation rate, the relatively high level of viability suggests that the inability of the double mutant to form binucleate cells is likely not due to a catastrophic defect in repairing Spo11-induced DSBs (see below).

A meiotic time course of WT, *nup2Δ, ndj1Δ and* double mutant combinations involving the truncations mutations described in Figure 2 revealed that only the *nup2Δ1-175* mutant gave delayed MI kinetics similar to *nup2Δ and* failed to progress beyond the mononucleate stage in the absence of Ndj1 (Figure 1 and Figure 4A). On the other hand, the *nup2-MAR ndj1Δ* double mutant gave WT MI division kinetics and failed to give a synthetic phenotype with *ndj1Δ*. This is another indication that the Nup2-MAR region alone is important for meiotic progression.

Since Csm4 is the likely KASH protein linking telomeres to cytoskeletal actin in budding yeast (Fridolfsson and Starr 2010, Wanat *et al*. 2008, Kosaka *et al*. 2008), we tested if the *csm4Δ* mutation in combination with *nup2Δ* would give a synthetic phenotype similar to *nup2Δ ndj1Δ.* First we looked at the sporulation efficiency of the single and double mutants and found that while the 92% of *nup2Δ and* 76% of *csm4Δ* cells produced two or more spores (n = 200), only 1% of *nup2Δ csm4Δ* double mutant cells (n = 200) produced spores, similar to what we observed for *nup2Δ ndj1Δ* (1.5% (n = 200; Table 1). Likewise, in a meiotic time-course experiment, nuclear divisions were nearly absence in the *nup2Δ csm4Δ* double mutant, which is similar to what we observed for *nup2Δ ndj1Δ* (Figure S4).

### Cause of elevated spore death in *nup2Δ*, *nup60Δ, htz1Δ, and ndj1Δ*

We recently developed a suite of R-scripts that gives a computational estimate of the rate of MI nondisjunction for a given strain based on the fractional incidence of 4, 3, 2, 1, and 0 viable spore tetrads (Chu and Burgess 2016). We applied this approach to infer the causes of spore death in WT, *ndj1Δ nup2Δ, nup60Δ, and htz1Δ* (Table 1). TetFit finds the best-fit values for calculated spore death due to MI-nondisjunction (NDD) and random spore death (RSD). The NDD/RSD ratios for our WT and *ndj1Δ* data sets were 2.0%/0.9% and 15.8%/2.7%, respectively. In both cases, the cause of spore death was skewed to NDD. This is consistent with NDD/RSD outcomes of TetFit analysis of previously published data sets for these genotypes (Wanat et al. 2008; Chu and Burgess, 2016), where the calculated NDD/RSD ratios for WT and *ndj1Δ* were 1.8%/1.8% and 20.8%/5.8%, respectively (Figure S5).

We next used TetFit to analyze the observed tetrad distributions of *nup2Δ, nup60Δ, and htz1Δ*. In all cases, these strains gave greater NDD compared to WT, suggesting that spore death in these mutants can be attributed, in part, to MI-ND. However, the best fit NDD/RSD ratios in these mutants were skewed to greater RSD values: *nup2Δ* was 5.2%/9.9%; *nup60Δ* was 8.9%/23.1%; and *htz1Δ* was 7.5%/29.4%. Thus, unlike *ndj1Δ*, the cause(s) of spore death in *nup2Δ, nup60Δ, and htz1Δ* appear to be predominantly due to RSD rather than MI-ND (Figure S5), suggesting these nuclear envelope-associated proteins act in a different pathway from those affected by Ndj1.

To analyze the confidence of the output values from TetFit, we ran 50 independent simulations of 4, 3, 2, 1, and 0 viable spore tetrads based on the calculated best fit NDD and RSD values found for each mutant. Simulated dissection data was then analyzed by TetFit.Test to estimate the best fitting NDD and RSD. The output of these 50 simulations of all five strains is shown in Figure 4B. All three annotated R-scripts can be found in the supplemental methods.

### Deletion of *SPO11* suppresses the block to nuclear divisions in *nup2Δ ndj1Δ*

One possible reason for the failure of the *nup2Δ ndj1Δ* double mutant to undergo nuclear division is that this strain also fails to progress past pachytene due to the activation of the recombination checkpoint. We thus tested if the block to nuclear divisions could by bypassed by also deleting *SPO11*.

Interestingly, while the *ndj1Δ nup2Δ spo11Δ* triple mutant could form binucleate cells at greater than 85% efficiency, it still gave delayed division kinetics that were indistinguishable from *nup2Δ* or even the *nup2Δ spo11Δ* double mutant (Figure 4C). Surprisingly, while the *nup2Δ and spo11Δ* single mutants sporulated at high efficiency (92.0% and 93.5%, respectively), the *nup2Δ spo11Δ* double mutant sporulated at very low levels (0.5%), suggesting a more complex genetic interaction than expected.

Finally, we tested if the delay phenotype of *nup2Δ* could be due to the activation of the DNA repair or the spindle assembly checkpoint and found that neither *rad9Δ* nor *mad3Δ* mutations relieved the delay conferred by *nup2Δ* (Figure S6; Cheslock *et al*. 2005; Cartagena-Lirola *et al*. 2008).

### The *nup2Δ ndj1Δ* mutant undergoes two rounds of SPB duplication without segregating chromosomes

The synthetic failure to form binucleate cells in the *nup2Δ ndj1Δ* double mutant provided a robust marker for further analyzing Nup2’s role in meiosis. To better understand this phenotype, we asked if the *nup2Δ ndj1Δ* double mutant cells had defects in duplicating and/or separating their SPBs. For this analysis we expressed the fluorescently-tagged SPB component Spc42-GFP in WT and mutant cells (Shirk *et al*. 2011). Expression and incorporation of Htb2-mRuby2 (a red fluorescently-tagged version of histone H2A) into chromatin allowed for us to also follow nuclear divisions. The most striking feature of this analysis was that two rounds of SPBs duplication and separation occurred, yet the cells remained mononucleate (Figure 5A). In the *nup2Δ ndj1Δ* mutant, we observed that 78% of cells gave ≥ 2 Spc42-GFP (n = 200) 12 hrs after transfer to SPM compared to 94% of WT and 90% of both *nup2Δ and ndj1Δ* single mutants. This is in stark contrast to the less than 5% of cells that were also binucleate. These results suggest that the *nup2Δ ndj1Δ* double mutant undergoes a metaphase I-like arrest without undergoing a cell cycle arrest.

**Figure 5.**
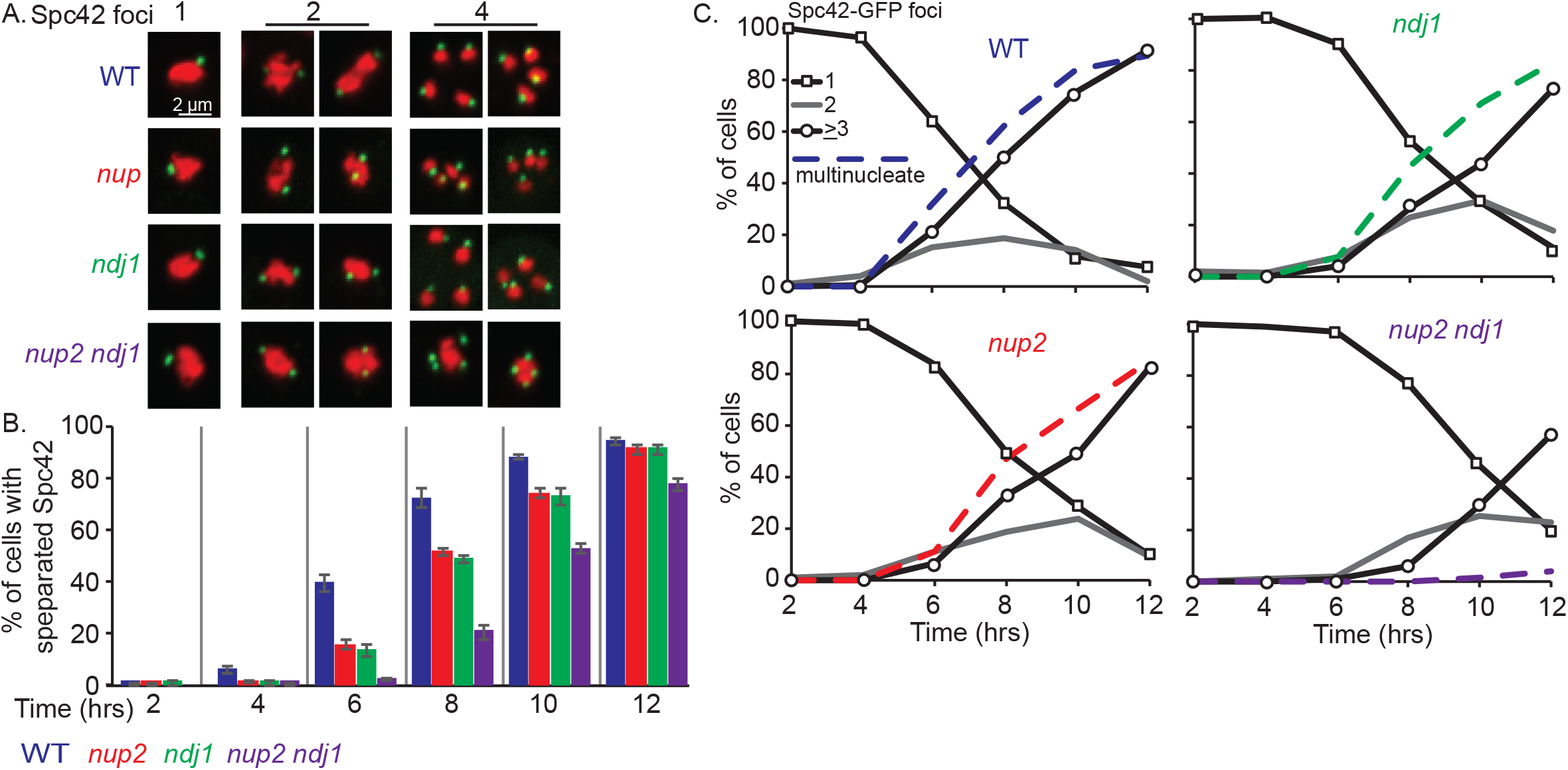
Meiotic time course to assay SPB duplication and separation in WT, *nup2Δ ndj1Δ*, and *nup2Δ ndj1Δ* cells. **A**. Representative images of with 1, 2, or 4 Spc42-GFP foci in fixed cells taken from a culture 12 hrs after transfer to SPM. Spc42-GFP is shown in green and Htb2-mRuby2 is shown in red. **B**. Kinetics of the percent of cells with separated Spc42-GFP foci in a meiotic time course at the indicated hrs after transfer to SPM. WT (Blue; SBY5963), *nup2Δ* (Red; SBY5969), *ndj1Δ* (Green; SBY5987), and *nup2Δ ndj1Δ* (Purple; SBY5993). **C**. Percentage of cells with 1 Spc42-GFP focus is shown in solid black line with open squares; cells with 2 Spc42-GFP foci are shown in gray; cells with ≥ 3 Spc42-GFP foci are shown black with open circles. Multinucleate cells were scored by DAPI staining of nuclei (dashed line).

Using a meiotic time-course assay, we visually assessed the number of cells with ≥ 2 Spc42-GFP foci in the four strains. Separation of Spc42-GFP foci in *nup2Δ and ndj1Δ* single mutants gave delayed kinetics compared to WT (Figure 5, B and C), which is not surprising given that they both show delays of meiotic prophase stages. This unusual phenotype allowed us to determine if the *nup2Δ and ndj1Δ* alleles show epistasis with respect to the timing of Spc42-GFP separation. We found that the time at which 50% of cells had duplicated Spc42-GFP foci was delayed ∼1 hour in the *nup2Δ* mutant and ∼1.3 hrs in the *ndj1Δ* mutant compared to WT, while the double mutant exhibited ∼2.9 hour delay (Figure 5C). This general lack of epistasis, compared to the other strong synthetic block to nuclear divisions, indicates that the primary defects in *nup2Δ and ndj1Δ* single mutants that lead to delayed SPB separation are due to defects in separate pathways. The severe synthetic phenotype of the double mutant thus appears to be neomorphic and may arise by the formation of a poisonous intermediate or product that depends on *SPO11* to form (Figure 4C).

### DSB formation and repair in *nup2Δ*, *ndj1Δ, and nup2Δ ndj1Δ*

We next tested if the molecular basis for the synthetic failure to form binucleate cells in *nup2Δ ndj1Δ* would be evident in one or more stages governing the repair of Spo11-induced DSBs. We thus measured recombination intermediates and products in WT, *nup2Δ, ndj1Δ, and nup2Δ ndj1Δ* using the *HIS4LEU2* DSB hotspot (Figure 6, A, B, and C; Storlazzi *et al*. 1995; Hunter and Kleckner 2001).

**Figure 6.**
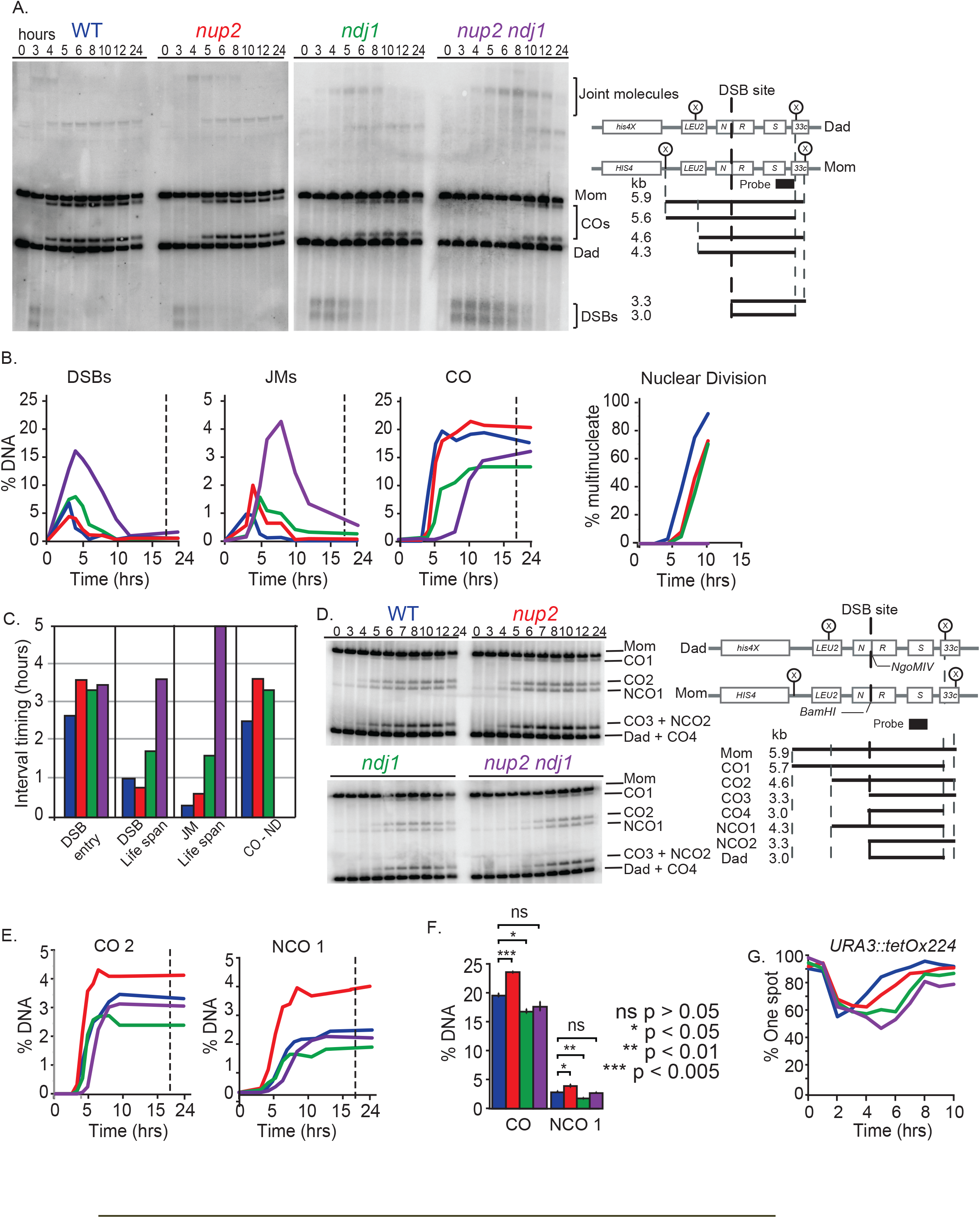
Time-course analysis of precursors, intermediates and products of homologous recombination at the *HIS4LEU2* hot spot in WT, *nup2Δ ndj1Δ*, and *nup2Δ ndj1Δ* cells. **A**. Representative Southern blots and schematic of the *HIS4LEU2* hotspot region showing the position of the Xho1 cut sites (x). The predicted sizes for the starting chromosomes from the *HIS4LEU2* strain (mom) and *his4XLEU2* strain (Dad), DSBs (3.3 and 3.0 kb), JMs, and CO products are shown. **B**. Amounts of DSB and recombinant products as a fraction of total DNA for each lane in a time course assay for WT (Blue; SBY1903), *nup2Δ* (Red; SBY3945), *ndj1Δ* (Green; 1904), and *nup2Δ ndj1Δ* (Purple; SBY3983). Right: Kinetics of forming multinucleate cells in the same time course. **C**. Lifespan analysis for DSB and JMs was performed as described in Figure 3. The interval between the time at which 50% of CO have formed and the time at which 50% of cells have undergone nuclear division is shown for all strains except *nup2Δ ndj1Δ*, which doesn’t undergo nuclear divisions. Lifespan analysis is based on maximum levels of DSBs measured for *sae2Δ*, *nup2Δ sae2Δ and* the *nup2Δ ndj1Δ sae2Δ* double mutant (Padmore *et al*. 1991). For comparison, lifespans for *ndj1Δ* were normalized to *sae2Δ* levels (Wu and Burgess 2006; Wanat *et al*. 2008) **D**. Southern blot analysis of CO and NCO formation in WT, *nup2Δ*, *ndj1Δ*, and *nup2Δ ndj1Δ*. The left side shows a Southern blot of a meiotic time course of WT (Blue; SBY5826), *nup2Δ* (Red; SBY5832), *ndj1Δ* (Green; SBY5838), and *nup2Δ ndj1Δ* (Purple; SBY5844). The right side shows the schematic and recombination products of the *HIS4LEU2* hotspot used. DNA was digested with XhoI and NgoMIV. **E**. CO2 and NCO1 quantifications as percent of total DNA. **F**. CO and NCO quantifications of WT (blue), *nup2Δ* (red), *ndj1Δ* (green), and *nup2Δ ndj1Δ* (purple) from Figure S7 A and B. Data are represented as mean of three independent replicates ± SD. P values are based on a t-test with multiple comparisons adjustment using the Holm method. Representative blots are shown in Figure S7. **G.** Pairing analysis of strains containing TetO arrays integrated at *URA3 and* expressing TetR-GFP fusion protein. Homologs were scored as paired if only a single GFP focus could be observed and unpaired if two GFP foci were observed. All strains are *ndt80Δ*. The pairing levels of WT (Blue; SBY5826), *nup2Δ* (Red; SBY5832), *ndj1Δ* (Green; SBY5838), and *nup2Δ ndj1Δ* (Purple; SBY5844) during a meiotic time course. 200 cells were scored for each time point.

**Figure 7.**
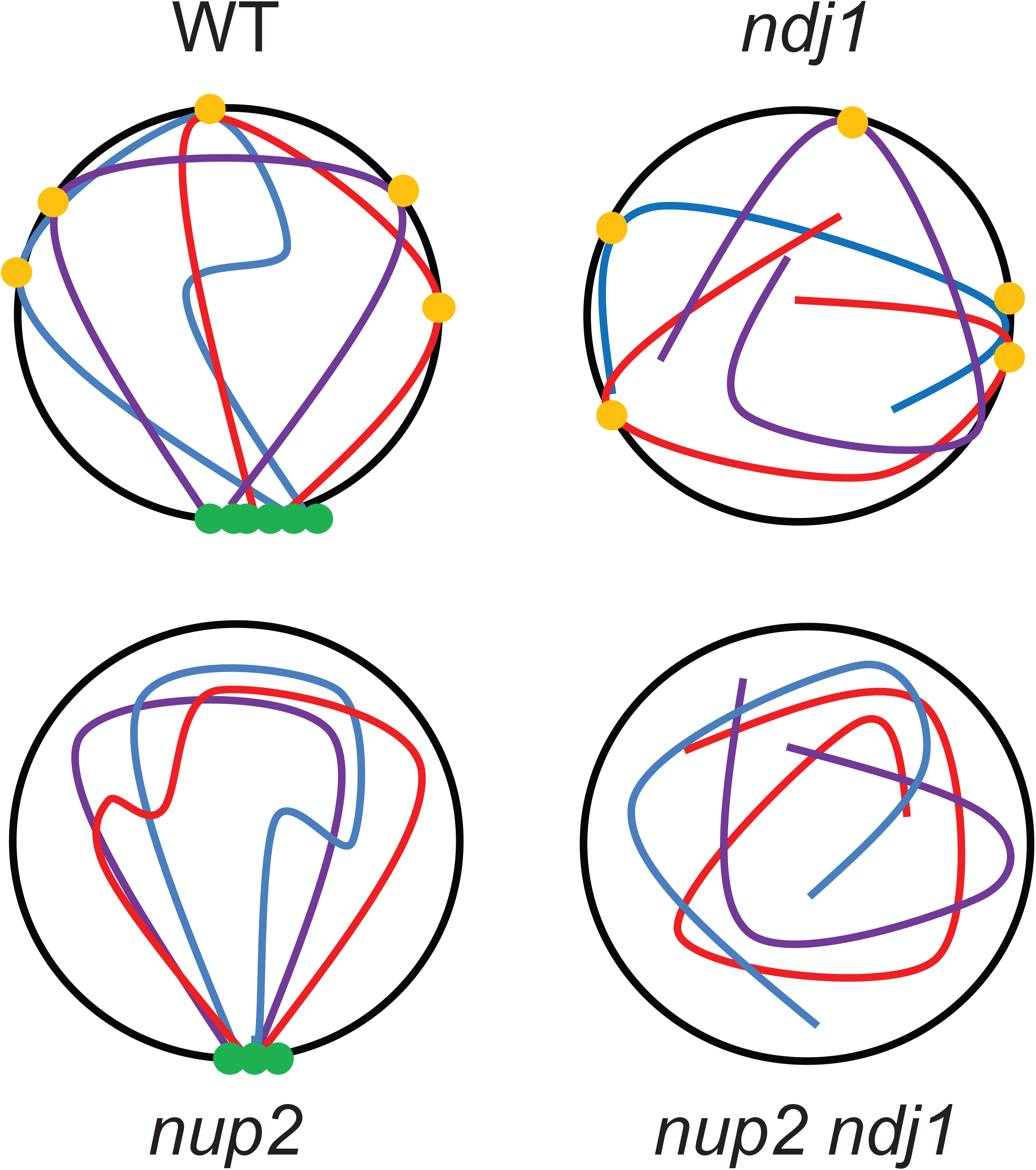
Model for two levels of chromosome organization in the meiotic nucleus supported by Ndj1 and Nup2. Loss of either pathway allows for meiotic progression, albeit with delay in nuclear division and progression through meiotic prophase. In the absence of both Ndj1 and Nup2, chromosome organization is impacted so that chromosomes are physically unable to separate. Another possibility is that the *nup2Δ* mutation impacts the ability of sister chromatids to separate
which is not mutually exclusive of the given model (not shown).

Using the total DSB levels found for *sae2Δ and nup2Δ sae2Δ* strains above, we calculated the interval timing between the formation and repair of DSBs was in WT and *nup2Δ and* found them to be similar, if not slightly shorter in *nup2Δ* (0.9 and 0.8 hrs, respectively). This was consistent with the slightly shorter zygotene window we observed. As seen previously (Wanat *et al*. 2008), we found the DSB lifespan in the *ndj1Δ* mutant was about twice as long as WT (1.7 hrs) and this transition in *nup2Δ ndj1Δ* was ∼ 3.6 hrs. By simply adding the two effects we would expect less than one hr delay in DSB turnover in the *nup2Δ ndj1Δ* mutant, yet the delay we found was nearly three hrs longer (Figure 6C). Considered together, these results point to a synthetic defect in the repair of DSBs in the double mutant as a possible contributing factor to the failure of this strain to form binucleate cells.

### DNA joint molecule (JM) turnover in *nup2Δ*, *ndj1Δ*, and *nup2Δ ndj1Δ*

DSBs are processed into JMs that include single-end invasions and double-Holiday Junction intermediates (Hunter 2015; Lam and Keeney 2015). Disruption of the recombination program can also result in more complex forms of JMs (Oh *et al*. 2007; Kaur *et al*. 2015; Tang *et al*. 2015). We found that the JM lifespan in *nup2Δ* (0.6 hrs) was longer than WT (0.3 hrs), but not to the extent of *ndj1Δ* (1.6 hrs; Figure 6C). The most dramatic effect on JM lifespan, however, was seen in *nup2Δ ndj1Δ* double mutant (5.0 hrs; Figure 6C). These results point to the accumulation of JMs as a possible contributing factor of the nuclear-division failure of the *nup2Δ ndj1Δ* mutant. Even though it appears that most of the JMs are turned over, less efficient repair at any given other locus could impede chromosome separation.

### CO formation to MI division timing is delayed in *nup2Δ and ndj1Δ* compared to WT

The timing of CO formation to MI provides an additional landmark to account for the post-DSB contribution to the *nup2Δ* delay. The *nup2Δ and ndj1Δ* mutants exhibited a 0.9 and 0.6 hr delay from 50% CO levels to the time at which 50% of cells underwent nuclear division, respectively, compared to WT (Figure 6C). The *nup2Δ ndj1Δ* mutant could not be analyzed in this way due to the failure to segregate chromosomes. These data, and those reported above, suggest that there are at least three stages of meiotic prophase that are extended in the *nup2Δ* mutant compared to WT (approximate values in hours): a period prior to or during DSB formation (+ 0.9), the lifespan of JMs (+ 0.3), and the interval between CO formation and chromosome separation at MI (+ 0.9; Figure 5C). The *ndj1Δ* mutant also exhibits a delay in DSB and JM turnover as reported previously (Wu and Burgess 2006; Wanat *et al*. 2008). These latter two intervals are greatly extended in *nup2Δ ndj1Δ* compared to WT (+ 2.7 and + 4.7 hrs, respectively; Figure 6C). Thus, by acting alone or with Ndj1, Nup2 appears to influence multiple steps of the meiotic program, from DSB formation to the separation of chromosomes at MI.

### CO and noncrossover (NCO) levels are elevated in *nup2Δ* compared to WT

In the time course study described above, the *nup2Δ* mutant gave slightly elevated CO levels compared to WT (Figure 6B). We followed up this observation by analyzing a modified *HIS4LEU2* allele, whereby cutting by both XhoI and NgoMIV can recover NCO products that have undergone gene conversion but not exchange of flanking markers (Figure 6D). Using this strain, in a time course assay, we found that CO and NCO levels of well separated products in *nup2Δ* were overall higher than in WT (Figure 6E). We next carried out a separate experiment done in triplicate using just XhoI to analyze total CO levels at 12 hrs after transfer to SPM. Accordingly, we found that the total CO levels in the *nup2Δ* mutant to be significantly higher than WT (23.7% vs. 19.6%, respectively, t-test, p = 0.004; Figure 6F and Figure S7A). Using XhoI and NgoMIV we found NCO levels to be slightly higher than WT: 3.3% vs. 2.4%, t-test, p = 0.04; Figure 6F and Figure S7B). Taken together, the increased total levels of DSBs and recombination products we see in the absence of Nup2 suggest that Nup2 may have a negative influence on regulating DSB formation (see discussion).

### Nup2 and Ndj1 contribute independently to the efficiency of homolog pairing

It is well established that homologous chromosome pairing is delayed in *ndj1Δ* (Chua and Roeder 1997; Peoples-Holst and Burgess 2005; Conrad *et al*. 2008; Lui *et al*. 2013). We used a previously described “one-spot, two-spot” assay (Brar *et al*. 2009; Lui *et al*. 2013) in which chromosome *V* homologs, each with an array of *tetO*-repeats inserted at *URA3* were visualized in individual cells expressing a TetR-GFP fusion protein. Binding of this protein to the *tetO*-repeats gives a one-focus “spot” if the loci are paired and two foci if they are unpaired (Michaelis *et al*. 1997). This experiment was done in an *ndt80Δ* mutant strain background where pairing persists since cells arrest in the pachytene stage. As seen previously, over 80% of cells gave a one-spot signal at the time of transfer to SPM (at the G0 stage; Figure 5E; Brar *et al*. 2009; Lui *et al*. 2013). At 2 hrs, WT, *nup2Δ*, *ndj1Δ*, and *nup2Δ ndj1Δ* strains exhibited timely disruption of G0 pairing, which coincides with DNA replication timing (Figure 5E; Brar *et al*. 2009). Pairing was re-established in both *ndj1Δ and nup2Δ* mutants, albeit with some delay compared to WT (Figure 6E). Pairing in *nup2Δ ndj1Δ* cells was further delayed compared to either single mutant suggesting that the contributions of Nup2 and Ndj1 to homolog pairing occur through independent pathways (Figure 6E).

## DISCUSSION

Here we show that a 125 amino-acid region of the full-length 720 amino acids of Nup2 is necessary and sufficient to carry out its meiotic role(s), thus behaving as a <u>m</u>eiotic <u>a</u>utonomous <u>r</u>egion (MAR). The MAR sequence alone is sufficient to confer localization to the nuclear periphery and has the same genetic dependency on Nup60 as full length Nup2. We predict that this MAR domain is responsible for Nup2’s localization to the nuclear periphery. An N-terminal fragment of Nup2 that includes the MAR was previously shown to complement the synthetic lethality between *nup2Δ and nup1-8* (Loeb *et al*. 1993). Therefore, the sequence is also likely to be functionally relevant for vegetative growth. There are two predicted SUMOylation motifs within the MAR at residues aa153 and aa170 (Ren *et al*. 2009; Zhao *et al*. 2014). SUMOylation has been shown to regulate several chromosome-based events of meiosis. Mutating these sites may provide further insight into Nup2’s meiotic function (Watts and Hoffmann 2011; Voelkel-Meiman *et al*. 2013).

There is compelling evidence to suggest that Nup2’s role in meiosis may be conserved. In mouse, mNup50 and its paralog, n50rel, are enriched in the testis (Fan *et al*. 1997; Smitherman *et al*. 2000; Park *et al*. 2016). Moreover, loss of mNup50 leads to the apoptosis of primordial germ cells resulting in an absence of germ cells (Park *et al*. 2016). Using a BLAST search across phyla, we found that only Nup50 homologs gave a significant score from a BLAST search with the MAR sequences (data not shown).

Several lines of evidence demonstrate a nucleoplasmic role for Nup2/hNup50 apart from the NPC. In *Drosophila melanogaster*, nucleoplasmic Nup50 binds to developmental and heat shock-induced puffs of salivary gland chromosomes in larva (Kalverda *et al*. 2010). In *Aspergillus nidulans,* Nup2 is an essential protein and requires its binding partner NupA to fulfill its functions. Moreover, Nup2 relocates from the NPC during interphase to chromatin during mitosis (Markossian *et al*. 2015). In mouse, mNup50 has been shown to play a role in regulating chromatin architecture, independent of transport function, and its association with mNup153 (yeast Nup60 homolog; Buchwalter *et al*. 2014). In this case, only the N-terminal half of hNup50 is required for mobility between the NPC and the nucleoplasm, which includes region defined here as the MAR. We show that the enrichment of the Nup2-MAR at the nuclear periphery depends on Nup60, however, *nup60Δ* mutants do not give the same synthetic meiotic block to nuclear division as occurs in the *nup2Δ ndj2Δ* double mutant.

In budding yeast, Nup2 (hNup50), Nup60 (hNup153) and Nup53 are conserved nonessential nups that are primarily found on the nuclear basket (Loeb *et al*. 1993; Fan *et al*. 1997; Hood *et al*. 2000). Nup2 and Nup60 are implicated in aspects of nuclear architecture that impact processes such as the DNA damage response, transcriptional silencing, boundary activity, transcriptional memory, mRNA processing, and mRNA export (Fahrenkrog *et al*. 2000; Denning *et al*. 2001; Bukata *et al*. 2013; Ptak *et al*. 2014). Additionally, Nup2 localizes to hundreds of chromosomal loci suggesting it may have a broad role in chromosome organization (Casolari *et al*. 2004; Dilworth *et al*. 2005; Schmid *et al*. 2006). It may be that one or more of these established roles are also required to facilitate normal progression through meiotic prophase.

Several aspects of our data analyzing single and double mutant phenotypes are consistent with the notion that Nup2 functions in a pathway independent of Ndj1 in meiosis. In addition to the block to nuclear division, we also observed synthetic defects in homologous recombination, an additive delay in homolog pairing and SPB separation, and different modalities of spore inviability.

The *nup2Δ ndj1Δ* synthetic phenotype is unusual since cells form four separate SPBs without undergoing nuclear division. This is similar to mutants disrupting Top3-Rmi1 decatenase, which fail to undergo nuclear division due to unresolved JMs without disrupting the meiotic program (Gangloff *et al*. 1999; Kaur *et al*. 2015; Tang *et al*. 2015), in a phenomenon known as “meiotic catastrophe” (Jessop and Lichten 2008; Oh *et al*. 2008). The *nup2Δ ndj1Δ* double mutant differs from the *top3/rmi1* mutants since the former fails to sporulate while latter continues the meiotic program through sporulation. A second way binucleate formation can be blocked is through the failure to lose connections between sister chromatids, as is seen in the absence Tid1, which causes abnormal persistence of the Mcd1 and Rec8 cohesin proteins (Kateneva *et al*. 2005). Nuclei in the *tid1Δ* mutant, however give a stretched, dumbbell-shaped appearance that we do not see in the *nup2Δ ndj1Δ* mutant. In each of the above cases, including *nup2Δ ndj1Δ*, deletion of *SPO11* was found to bypass the block to nuclear divisions Thus from a first appearance, it would appear that the inability of the *nup2Δ ndj1Δ* mutant to undergo nuclear divisions is due to the presence of an aberrant intermediate(s) or products of homologous recombination.

One key difference from the *top3/rmi1 and tid1* mutants is that deleting *SPO11* in the *nup2Δ ndj1Δ* background, or even *nup2Δ* alone, does not to suppress the delay phenotype seen when *NUP2* is absent. Moreover, the unexpected sporulation defect in the *nup2Δ spo11Δ* double mutant suggests a more complex relationship between the two mutations that may impinge on the ability of sister chromatids to separate from one another at meiosis I anaphase, albeit without forming dumbbells. If this were the case, then the formation of even normal CO products would prevent nuclear division, which is seen when either separase (Esp1) or cleavage of the Rec8 cohesin subunit is inactivated (Buonomo *et al*. 2000) or the inability to form monopolar kinetochore attachments in meiosis I (TÓth *et al*. 2000).

The synthetic phenotype of *nup2Δ spo11Δ* is relevant since Spo11, in addition to its role in forming DSBs, plays an earlier role in establishing chromosome axis structure during meiotic S-phase and promoting the recruitment of chromosome accessory factors that might be sensitized in the absence or Nup2 (Cha *et al*. 2000; Kugou *et al*. 2009; Panizza *et al*. 2011). Notably, the increased levels of DSB and recombination products (both CO and NCO) we see in the *nup2Δ* mutant are similar to other mutants that show misregulation of DSB formation in the context of aberrant axis composition. For example, there is strong evidence that a Mec1/Tel1-dependent pathway actively suppresses DSB formation by phosphorylation of Spo11 interacting partner Rec114 at the chromosome axis (Zhang *et al*. 2011; Carballo *et al*. 2013). It is possible that Nup2 could influence this pathway, which is readily testable.

Finally, Kleckner and colleagues have pointed out several lines of evidence to suggest that nondisjunction in the *ndj1Δ and csm4Δ* mutants may be due to defects in the relationship between sister chromatids (Wanat *et al*. 2008). We also report here that deletion of *CSM4* also gives a synthetic meiotic defect by blocking both nuclear division and sporulation in the absence of Nup2. Thus, multiple observations point to an early role for Nup2 function in establishing a normal chromosome axis structure that underlies nearly every step of meiotic chromosome dynamics. Experiments designed to test these possibilities are in progress.

The region near the inner-nuclear membrane is enriched for specific DNA sequences, macromolecular complexes including the NPC and the SPB, and chromatin silencing factors (Taddei and Gasser 2012). The inner-nuclear membrane also plays a special role in meiosis by tethering telomeres to SUN/KASH bridges that spans the inner and outer nuclear membrane. These connections are the basis of dramatic telomere led motion that occurs during meiotic prophase (Koszul *et al*. 2008; Fridolfsson and Starr 2010). We propose that the higher order organization of chromosomes in the nucleus, mediated by Nup2, acts in parallel with Ndj1-mediated functions to support the dynamic chromosome interactions associated with homolog pairing, synapsis, and homologous recombination. Disruption of either configuration alone can be accommodated to complete meiosis, albeit with some delay, however, when both are disrupted, aberrant inter-sister and/or interhomolog interactions prevents their separation at anaphase I (Figure 6). This may explain why Nup2 is largely dispensable for normal vegetative growth under non-stressed conditions where the dynamic aspects of meiotic chromosome behavior are not present.

## Acknowledgements

We would like to thank Dan Starr, JoAnne Engebrecht, Anne Britt, James McGehee, and An Nguyen for their insightful review of this manuscript. We thank Neil Hunter for yeast strains. We thank the Kaback lab for the ZIP1-GFP plasmid. We thank Akira Shinohara for sharing detailed protocols. The authors have declared that no conflicting interests exist. This work was funded by NIH R01 GM075119 award to S.M.B and pilot funds from the UC Davis Proteomics Core.

## Abbreviations List

WT: Wild type
CO: Crossover
DSB: Double-strand break
JM: Joint molecule
MI: Meiosis I division
NCO: Noncrossover
NPC: Nuclear pore complex
Nup: Nucleoporin
SC: Synaptonemal complex
SPB: Spindle-Pole Body

## Summary of supplemental material

**File S1** Excel spread sheet of mass spectrometry analysis.

**File S2** Background and annotated R-scripts with instructions for using TetSim, TetFit, and TetFit.Test.

**Figure S1 Time course analysis to measure the kinetics of nuclear divisions in WT, *rif1Δ, rif2Δ, rif1Δ rif2Δ*** Time course analysis was done as in Figure 1: WT (SBY1903), *rif1Δ* (SBY3948), *rif2Δ* (SBY 4928), and *rif1Δ rif2Δ* (SBY5997).

**Figure S2 Colony growth of nup mutants on YPD and minimal media at different temperatures. A**. rowth of WT and *nup* mutants at 20° C, 25° C, 30° C, and 37° C on YPD for 48 and 72 hrs. **B**. Growth of WT and *nup* mutants at 20° C, 25° C, 30° C, and 37° C on minimal media for 72 or 120 hrs.

**Figure S3 Localization of Srp1-GFP and Cse1-GFP in nup2 mutants** Representative maximum intensity Z-projections of fixed WT (top), *nup2Δ* (middle), and *nup2-MAR* cells (bottom) expressing Srp1-GFP (left) or Cse1-GFP (right) taken from an exponentially dividing culture.

**Figure S4 Time course analysis to measure the kinetics of nuclear divisions in WT, *nup2Δ, csm4Δ, nup2Δ csm4Δ*** Time course analysis was done as in Figure 1: WT (SBY1903), *nup2Δ* (SBY3948), *csm4Δ* (SBY 4040), and *nup2Δ csm4Δ* (SBY4045).

**Figure S5 Kinetics of nuclear divisions in Outcome of TetSim and TetFit on tetrad data from WT*, nup2Δ, ndj1Δ, nup60Δ*, and *htz1Δ.* A**. Use of TetSim to compare of observed spore death with a simulated distribution based on random spore death (RSD). TetSim simulates a distribution of 4:0, 3:1, 2:2, 1:3, and 0:4 tetrads based on the observed fraction of viable cells (Chu and Burgess 2016) and compares the 99% cutoff values for the simulated distribution to the observed distribution for each genotype. The number of simulations for each panel is equal to the number of tetrads dissected experimentally. A distribution of dissected tetrad simulations is shown (in red, 4:0; green, 3:1; blue, 2:2; teal, 1:3: and purple, 0:4) with the upper and lower dashed lines represent 99% and 1% percentiles, respectively. The observed frequencies of live:dead tetrad distributions for individual strains are shown as solid lines. Comparison of simulated tetrad distributions with mutant data from previously published data sets for WT and *ndj1Δ* (Wanat *et al*. 2008). Note the fraction of viable spore in the *ndj1Δ* mutant was higher in our study than previously reported by Wanat et a., 2008. **B**. Best fitting distributions of 4 (red), 3 (green), 2 (blue), 1 (teal), and 0 (purple)-viable spore tetrads in WT, *ndj1Δ*, *nup2Δ*, *nup60Δ*, and *htz1Δ*, mutants generated by TetFit (Chu and Burgess, 2016). Analysis of published data in Wanat et al., 2008 was also included in the analysis as a control. The observed tetrad distributions of genotype (O, middle bar), the expected tetrad distributions with the best fitting MI-ND and RSD (E, right bar), and the expected tetrad distributions using the calculated WT MI-nondisjunction rate and the best fitting RSD (W, left bar). Significance is noted as * *P* < 0.05; *** *P* < 0.001 (chi-squared, Holm corrected). The calculated values for NDD, RSD and the NDD/RSD ratio are shown below each bar plot for each mutant strain analyzed.

**Figure S6 The impact of *nup2Δ* on delaying nuclear divisions is not suppressed by *rad9Δ* or *mad3Δ*** Left: Time course experiment as described in Figure 1. WT (Blue), *nup2Δ* (Red), *rad9Δ* (Green), and *nup2Δ rad9Δ* (Purple). Right: Same except for *mad3Δ and nup2Δ mad3Δ*.

**Figure S7 Analysis of terminal CO and NCO levels at *HIS4LEU2* in WT, *nup2Δ*, *ndj1Δ*, and *nup2Δ ndj1Δ.* A**. Representative Southern blot for CO analysis. The right side shows the schematic of relevant markers at the *HIS4LEU2* hotspot and the location of the DNA probe. Isolated genomic DNA was digested with XhoI prior to gel electrophoresis. The size of expected bands is also indicated. **B**. Representative Southern blot for NCO analysis. Same as in part A although DNA was digested with XhoI and NgoMIV.

## References

Alani E., R. Padmore and N. Kleckner, 1990 Analysis of wild-type and rad50 mutants of yeast suggests an intimate relationship between meiotic chromosome synapsis and recombination. Cell 61: 419–436.

Benjamini, Y., and Y. Hochberg, 1995 Controlling the false discovery rate: a practical and powerful approach to multiple testing. Journal of the Royal Statistical Society. Series B (Methodological): 289–300.

Bishop, D. K., D. Park, L. Xu and N. Kleckner, 1992 DMC1: a meiosis-specific yeast homolog of E. coli recA required for recombination, synaptonemal complex formation, and cell cycle progression. Cell 69: 439–456.

Blitzblau, H. G., and A. Hochwagen, 2013 ATR/Mec1 prevents lethal meiotic recombination initiation on partially replicated chromosomes in budding yeast. Elife 2: e00844.

Booth, J. W., K. D. Belanger, M. I. Sannella and L. I. Davis, 1999 The yeast nucleoporin Nup2p is involved in nuclear export of importin alpha/Srp1p. J Biol Chem 274: 32360–32367.

Borner, G. V., N. Kleckner and N. Hunter, 2004 Crossover/noncrossover differentiation, synaptonemal complex formation, and regulatory surveillance at the leptotene/zygotene transition of meiosis. Cell 117: 29–45.

Brar, G. A., A. Hochwagen, L. S. Ee and A. Amon, 2009 The multiple roles of cohesin in meiotic chromosome morphogenesis and pairing. Mol Biol Cell 20: 1030–1047.

Buchwalter, A. L., Y. Liang and M. W. Hetzer, 2014 Nup50 is required for cell differentiation and exhibits transcription-dependent dynamics. Mol Biol Cell 25: 2472–2484.

Bukata, L., S. L. Parker and M. A. D’Angelo, 2013 Nuclear pore complexes in the maintenance of genome integrity. Current opinion in cell biology 25: 378–386.

Buonomo, S. B., R. K. Clyne, J. Fuchs, J. Loidl, F. Uhlmann, et al., 2000 Disjunction of homologous chromosomes in meiosis I depends on proteolytic cleavage of the meiotic cohesin Rec8 by separin. Cell 103: 387–398.

Cao, L., E. Alani and N. Kleckner, 1990 A pathway for generation and processing of double-strand breaks during meiotic recombination in S. cerevisiae. Cell 61: 1089–1101.

Carballo, J. A., S. Panizza, M. E. Serrentino, A. L. Johnson, M. Geymonat, et al., 2013 Budding yeast ATM/ATR control meiotic double-strand break (DSB) levels by down-regulating Rec114, an essential component of the DSB-machinery. PLoS Genet 9: e1003545.

Cartagena-Lirola, H., I. Guerini, N. Manfrini, G. Lucchini and M. P. Longhese, 2008 Role of the Saccharomyces cerevisiae Rad53 checkpoint kinase in signaling double-strand breaks during the meiotic cell cycle. Molecular and cellular biology 28: 4480–4493.

Casolari, J. M., C. R. Brown, S. Komili, J. West, H. Hieronymus, et al., 2004 Genome-wide localization of the nuclear transport machinery couples transcriptional status and nuclear organization. Cell 117: 427–439.

Cha, R. S., B. M. Weiner, S. Keeney, J. Dekker and N. Kleckner, 2000 Progression of meiotic DNA replication is modulated by interchromosomal interaction proteins, negatively by Spo11p and positively by Rec8p. Genes Dev 14: 493–503.

Cheslock, P. S., B. J. Kemp, R. M. Boumil and D. S. Dawson, 2005 The roles of MAD1, MAD2 and MAD3 in meiotic progression and the segregation of nonexchange chromosomes. Nat Genet 37: 756–760.

Chikashige, Y., D. Q. Ding, H. Funabiki, T. Haraguchi, S. Mashiko, et al., 1994 Telomere-led premeiotic chromosome movement in fission yeast. Science 264: 270–273.

Chu, D. B., and S. M. Burgess, 2016 A computational approach to estimating nondisjunction frequency in Saccharomyces cerevisiae. G3: Genes| Genomes| Genetics: g3. 115.024380.

Chu, S., and I. Herskowitz, 1998 Gametogenesis in yeast is regulated by a transcriptional cascade dependent on Ndt80. Molecular cell 1: 685–696.

Chua, P. R., and G. S. Roeder, 1997 Tam1, a telomere-associated meiotic protein, functions in chromosome synapsis and crossover interference. Genes Dev 11: 1786–1800.

Conrad, M. N., A. M. Dominguez and M. E. Dresser, 1997 Ndj1p, a meiotic telomere protein required for normal chromosome synapsis and segregation in yeast. Science 276: 1252–1255.

Conrad, M. N., C. Y. Lee, G. Chao, M. Shinohara, H. Kosaka, et al., 2008 Rapid telomere movement in meiotic prophase is promoted by NDJ1, MPS3, and CSM4 and is modulated by recombination. Cell 133: 1175–1187.

Conrad, M. N., C. Y. Lee, J. L. Wilkerson and M. E. Dresser, 2007 MPS3 mediates meiotic bouquet formation in Saccharomyces cerevisiae. Proc Natl Acad Sci U S A 104: 8863–8868.

Cuperus, G., and D. Shore, 2002 Restoration of silencing in Saccharomyces cerevisiae by tethering of a novel Sir2-interacting protein, Esc8. Genetics 162: 633–645.

Danilevich, V. N., and E. V. Grishin, 2002 A New Approach to the Isolation of Genomic DNA from Yeast and Fungi: Preparation of DNA-containing Cell Envelopes and Their Use in PCR. Russian Journal of Bioorganic Chemistry 28: 136–146.

Denning, D., B. Mykytka, N. P. Allen, L. Huang, B. Al, et al., 2001 The nucleoporin Nup60p functions as a Gsp1p-GTP-sensitive tether for Nup2p at the nuclear pore complex. J Cell Biol 154: 937–950.

Dilworth, D. J., A. J. Tackett, R. S. Rogers, E. C. Yi, R. H. Christmas, et al., 2005 The mobile nucleoporin Nup2p and chromatin-bound Prp20p function in endogenous NPC-mediated transcriptional control. J Cell Biol 171: 955–965.

Dresser, M. E., 2009 Time-lapse fluorescence microscopy of Saccharomyces cerevisiae in meiosis. Methods Mol Biol 558: 65–79.

Fahrenkrog, B., W. Hubner, A. Mandinova, N. Pante, W. Keller, et al., 2000 The yeast nucleoporin Nup53p specifically interacts with Nic96p and is directly involved in nuclear protein import. Mol Biol Cell 11: 3885–3896.

Fan, F., C. P. Liu, O. Korobova, C. Heyting, H. H. Offenberg, et al., 1997 cDNA cloning and characterization of Npap60: a novel rat nuclear pore-associated protein with an unusual subcellular localization during male germ cell differentiation. Genomics 40: 444–453.

Finn, R. D., J. Mistry, J. Tate, P. Coggill, A. Heger, et al., 2010 The Pfam protein families database. Nucleic Acids Res 38: D211–222.

Fridolfsson, H. N., and D. A. Starr, 2010 Kinesin-1 and dynein at the nuclear envelope mediate the bidirectional migrations of nuclei. The Journal of cell biology 191: 115–128.

Gangloff, S., B. de Massy, L. Arthur, R. Rothstein and F. Fabre, 1999 The essential role of yeast topoisomerase III in meiosis depends on recombination. The EMBO Journal 18: 1701–1711.

Goldstein, A. L., and J. H. McCusker, 1999 Three new dominant drug resistance cassettes for gene disruption in Saccharomyces cerevisiae. Yeast 15: 1541–1553.

Hassold, T., and P. Hunt, 2001 To err (meiotically) is human: the genesis of human aneuploidy. Nat Rev Genet 2: 280–291.

Ho, H. C., and S. M. Burgess, 2011 Pch2 acts through Xrs2 and Tel1/ATM to modulate interhomolog bias and checkpoint function during meiosis. PLoS Genet 7: e1002351.

Holm, S., 1979 A simple sequentially rejective multiple test procedure. Scandinavian journal of statistics: 65–70.

Hood, J. K., J. M. Casolari and P. A. Silver, 2000 Nup2p is located on the nuclear side of the nuclear pore complex and coordinates Srp1p/importin-alpha export. J Cell Sci 113 (Pt 8): 1471–1480.

Horn, H. F., D. I. Kim, G. D. Wright, E. S. Wong, C. L. Stewart, et al., 2013 A mammalian KASH domain protein coupling meiotic chromosomes to the cytoskeleton. J Cell Biol 202: 1023–1039.

Huh, W. K., J. V. Falvo, L. C. Gerke, A. S. Carroll, R. W. Howson, et al., 2003 Global analysis of protein localization in budding yeast. Nature 425: 686–691.

Hunter, N., 2015 Meiotic recombination: the essence of heredity. Cold Spring Harbor perspectives in biology 7: a016618.

Hunter, N., and N. Kleckner, 2001 The single-end invasion: an asymmetric intermediate at the double-strand break to double-holliday junction transition of meiotic recombination. Cell 106: 59–70.

Ishida, T., and K. Kinoshita, 2008 Prediction of disordered regions in proteins based on the meta approach. Bioinformatics 24: 1344–1348.

Jessop, L., and M. Lichten, 2008 Mus81/Mms4 endonuclease and Sgs1 helicase collaborate to ensure proper recombination intermediate metabolism during meiosis. Molecular cell 31: 313–323.

Kalverda, B., H. Pickersgill, V. V. Shloma and M. Fornerod, 2010 Nucleoporins directly stimulate expression of developmental and cell-cycle genes inside the nucleoplasm. Cell 140: 360–371.

Kateneva, A. V., A. A. Konovchenko, V. Guacci and M. E. Dresser, 2005 Recombination protein Tid1p controls resolution of cohesin-dependent linkages in meiosis in Saccharomyces cerevisiae. The Journal of cell biology 171: 241–253.

Kaur, H., A. De Muyt and M. Lichten, 2015 Top3-Rmi1 DNA single-strand decatenase is integral to the formation and resolution of meiotic recombination intermediates. Molecular cell 57: 583–594.

Keeney, S., C. N. Giroux and N. Kleckner, 1997 Meiosis-specific DNA double-strand breaks are catalyzed by Spo11, a member of a widely conserved protein family. Cell 88: 375–384.

Kleckner, N., 2006 Chiasma formation: chromatin/axis interplay and the role(s) of the synaptonemal complex. Chromosoma 115: 175–194.

Kleckner, N., D. Zickler, G. H. Jones, J. Dekker, R. Padmore, et al., 2004 A mechanical basis for chromosome function. Proc Natl Acad Sci U S A 101: 12592–12597.

Kosaka, H., M. Shinohara and A. Shinohara, 2008 Csm4-dependent telomere movement on nuclear envelope promotes meiotic recombination. PLoS Genet 4: e1000196.

Koszul, R., K. P. Kim, M. Prentiss, N. Kleckner and S. Kameoka, 2008 Meiotic chromosomes move by linkage to dynamic actin cables with transduction of force through the nuclear envelope. Cell 133: 1188–1201.

Koszul, R., and N. Kleckner, 2009 Dynamic chromosome movements during meiosis: a way to eliminate unwanted connections? Trends Cell Biol 19: 716–724.

Kugou, K., T. Fukuda, S. Yamada, M. Ito, H. Sasanuma, et al., 2009 Rec8 guides canonical Spo11 distribution along yeast meiotic chromosomes. Molecular biology of the cell 20: 3064–3076.

Lafontaine, D. L., and D. Tollervey, 2000 Synthesis and assembly of the box C+ D small nucleolar RNPs. Molecular and Cellular Biology 20: 2650–2659.

Lam, I., and S. Keeney, 2015 Mechanism and regulation of meiotic recombination initiation. Cold Spring Harbor perspectives in biology 7: a016634.

Lee, C. Y., M. N. Conrad and M. E. Dresser, 2012 Meiotic chromosome pairing is promoted by telomere-led chromosome movements independent of bouquet formation. PLoS Genet 8: e1002730.

Lee, S., W. A. Lim and K. S. Thorn, 2013 Improved blue, green, and red fluorescent protein tagging vectors for S. cerevisiae. PLoS One 8: e67902.

Light, W. H., D. G. Brickner, V. R. Brand and J. H. Brickner, 2010 Interaction of a DNA zip code with the nuclear pore complex promotes H2A. Z incorporation and INO1 transcriptional memory. Molecular cell 40: 112–125.

Lim, H. H., T. Zhang and U. Surana, 2009 Regulation of centrosome separation in yeast and vertebrates: common threads. Trends in cell biology 19: 325–333.

Loeb, J., L. Davis and G. Fink, 1993 NUP2, a novel yeast nucleoporin, has functional overlap with other proteins of the nuclear pore complex. Molecular biology of the cell 4: 209–222.

Longtine, M. S., A. McKenzie, 3rd, D. J. Demarini, N. G. Shah, A. Wach, et al., 1998 Additional modules for versatile and economical PCR-based gene deletion and modification in Saccharomyces cerevisiae. Yeast 14: 953–961.

Lui, D. Y., C. K. Cahoon and S. M. Burgess, 2013 Multiple opposing constraints govern chromosome interactions during meiosis. PLoS Genet 9: e1003197.

Markossian, S., S. Suresh, A. H. Osmani and S. A. Osmani, 2015 Nup2 requires a highly divergent partner, NupA, to fulfill functions at nuclear pore complexes and the mitotic chromatin region. Mol Biol Cell 26: 605–621.

Matsuura, Y., A. Lange, M. T. Harreman, A. H. Corbett and M. Stewart, 2003 Structural basis for Nup2p function in cargo release and karyopherin recycling in nuclear import. EMBO J 22: 5358–5369.

McKee, A. H., and N. Kleckner, 1997 A general method for identifying recessive diploid-specific mutations in Saccharomyces cerevisiae, its application to the isolation of mutants blocked at intermediate stages of meiotic prophase and characterization of a new gene SAE2. Genetics 146: 797–816.

Michaelis, C., R. Ciosk and K. Nasmyth, 1997 Cohesins: chromosomal proteins that prevent premature separation of sister chromatids. Cell 91: 35–45.

Moens, P. B., and E. Rapport, 1971 Spindles, spindle plaques, and meiosis in the yeast Saccharomyces cerevisiae (Hansen). The Journal of Cell Biology 50: 344–361.

Oh, S. D., L. Jessop, J. P. Lao, T. Allers, M. Lichten, et al., 2009 Stabilization and electrophoretic analysis of meiotic recombination intermediates in Saccharomyces cerevisiae. Methods Mol Biol 557: 209–234.

Oh, S. D., J. P. Lao, P. Y.-H. Hwang, A. F. Taylor, G. R. Smith, et al., 2007 BLM ortholog, Sgs1, prevents aberrant crossing-over by suppressing formation of multichromatid joint molecules. Cell 130: 259–272.

Oh, S. D., J. P. Lao, A. F. Taylor, G. R. Smith and N. Hunter, 2008 RecQ helicase, Sgs1, and XPF family endonuclease, Mus81-Mms4, resolve aberrant joint molecules during meiotic recombination. Molecular cell 31: 324–336.

Padmore, R., L. Cao and N. Kleckner, 1991 Temporal comparison of recombination and synaptonemal complex formation during meiosis in S. cerevisiae. Cell 66: 1239–1256.

Panizza, S., M. A. Mendoza, M. Berlinger, L. Huang, A. Nicolas, et al., 2011 Spo11-accessory proteins link double-strand break sites to the chromosome axis in early meiotic recombination. Cell 146: 372–383.

Park, E., B. Lee, B. E. Clurman and K. Lee, 2016 NUP50 is necessary for the survival of primordial germ cells in mouse embryos. Reproduction 151: 51–58.

Peoples-Holst, T. L., and S. M. Burgess, 2005 Multiple branches of the meiotic recombination pathway contribute independently to homolog pairing and stable juxtaposition during meiosis in budding yeast. Genes Dev 19: 863–874.

Ptak, C., J. D. Aitchison and R. W. Wozniak, 2014 The multifunctional nuclear pore complex: a platform for controlling gene expression. Curr Opin Cell Biol 28: 46–53.

Puig, O., F. Caspary, G. Rigaut, B. Rutz, E. Bouveret, et al., 2001 The tandem affinity purification (TAP) method: a general procedure of protein complex purification. Methods 24: 218–229.

Ren, J., X. Gao, C. Jin, M. Zhu, X. Wang, et al., 2009 Systematic study of protein sumoylation: Development of a site-specific predictor of SUMOsp 2.0. Proteomics 9: 3409–3412.

Rigaut, G., A. Shevchenko, B. Rutz, M. Wilm, M. Mann, et al., 1999 A generic protein purification method for protein complex characterization and proteome exploration. Nature biotechnology 17: 1030–1032.

Rockmill, B., and G. S. Roeder, 1998 Telomere-mediated chromosome pairing during meiosis in budding yeast. Genes & development 12: 2574–2586.

Rothstein, R., 1991 Targeting, disruption, replacement, and allele rescue: integrative DNA transformation in yeast. Methods Enzymol 194: 281–301.

Scherthan, H., H. Wang, C. Adelfalk, E. J. White, C. Cowan, et al., 2007 Chromosome mobility during meiotic prophase in Saccharomyces cerevisiae. Proc Natl Acad Sci U S A 104: 16934–16939.

Schmid, M., G. Arib, C. Laemmli, J. Nishikawa, T. Durussel, et al., 2006 Nup-PI: the nucleopore-promoter interaction of genes in yeast. Mol Cell 21: 379–391.

Sheehan, M. J., and W. P. Pawlowski, 2009 Live imaging of rapid chromosome movements in meiotic prophase I in maize. Proc Natl Acad Sci U S A 106: 20989–20994.

Sheff, M. A., and K. S. Thorn, 2004 Optimized cassettes for fluorescent protein tagging in Saccharomyces cerevisiae. Yeast 21: 661–670.

Shirk, K., H. Jin, T. H. Giddings, M. Winey and H.-G. Yu, 2011 The Aurora kinase Ipl1 is necessary for spindle pole body cohesion during budding yeast meiosis. Journal of cell science 124: 2891–2896.

Shulga, N., N. Mosammaparast, R. Wozniak and D. S. Goldfarb, 2000 Yeast nucleoporins involved in passive nuclear envelope permeability. J Cell Biol 149: 1027–1038.

Sigrist, C. J., L. Cerutti, E. de Castro, P. S. Langendijk-Genevaux, V. Bulliard, et al., 2010 PROSITE, a protein domain database for functional characterization and annotation. Nucleic Acids Res 38: D161–166.

Smitherman, M., K. Lee, J. Swanger, R. Kapur and B. E. Clurman, 2000 Characterization and targeted disruption of murine Nup50, a p27(Kip1)-interacting component of the nuclear pore complex. Mol Cell Biol 20: 5631–5642.

Solsbacher, J., P. Maurer, F. Vogel and G. Schlenstedt, 2000 Nup2p, a yeast nucleoporin, functions in bidirectional transport of importin alpha. Mol Cell Biol 20: 8468–8479.

Sosa, B. A., A. Rothballer, U. Kutay and T. U. Schwartz, 2012 LINC complexes form by binding of three KASH peptides to domain interfaces of trimeric SUN proteins. Cell 149: 1035–1047.

Storlazzi, A., L. Xu, L. Cao and N. Kleckner, 1995 Crossover and noncrossover recombination during meiosis: timing and pathway relationships. Proc Natl Acad Sci U S A 92: 8512–8516.

Sym, M., J. A. Engebrecht and G. S. Roeder, 1993 ZIP1 is a synaptonemal complex protein required for meiotic chromosome synapsis. Cell 72: 365–378.

Taddei, A., and S. M. Gasser, 2012 Structure and function in the budding yeast nucleus. Genetics 192: 107–129.

Tang, S., M. K. Y. Wu, R. Zhang and N. Hunter, 2015 Pervasive and essential roles of the Top3-Rmi1 decatenase orchestrate recombination and facilitate chromosome segregation in meiosis. Molecular cell 57: 607–621.

Tapley, E. C., and D. A. Starr, 2013 Connecting the nucleus to the cytoskeleton by SUN-KASH bridges across the nuclear envelope. Curr Opin Cell Biol 25: 57–62.

Teixeira, M. T., M. Arneric, P. Sperisen and J. Lingner, 2004 Telomere length homeostasis is achieved via a switch between telomerase-extendible and -nonextendible states. Cell 117: 323–335.

Tóth, A., K. P. Rabitsch, M. Gálová, A. Schleiffer, S. B. Buonomo, et al., 2000 Functional genomics identifies monopolin: a kinetochore protein required for segregation of homologs during meiosis I. Cell 103: 1155–1168.

Ubersax, J. A., E. L. Woodbury, P. N. Quang, M. Paraz, J. D. Blethrow, et al., 2003 Targets of the cyclin-dependent kinase Cdk1. Nature 425: 859–864.

Voelkel-Meiman, K., L. F. Taylor, P. Mukherjee, N. Humphryes, H. Tsubouchi, et al., 2013 SUMO localizes to the central element of synaptonemal complex and is required for the full synapsis of meiotic chromosomes in budding yeast. PLoS Genet 9: e1003837.

Wanat, J. J., K. P. Kim, R. Koszul, S. Zanders, B. Weiner, et al., 2008 Csm4, in collaboration with Ndj1, mediates telomere-led chromosome dynamics and recombination during yeast meiosis. PLoS Genet 4: e1000188.

Watts, F. Z., and E. Hoffmann, 2011 SUMO meets meiosis: an encounter at the synaptonemal complex: SUMO chains and sumoylated proteins suggest that heterogeneous and complex interactions lie at the centre of the synaptonemal complex. Bioessays 33: 529–537.

Wotton, D., and D. Shore, 1997 A novel Rap1p-interacting factor, Rif2p, cooperates with Rif1p to regulate telomere length in Saccharomyces cerevisiae. Genes & development 11: 748-760.

Wu, H. Y., and S. M. Burgess, 2006 Ndj1, a telomere-associated protein, promotes meiotic recombination in budding yeast. Mol Cell Biol 26: 3683–3694.

Wu, H. Y., H. C. Ho and S. M. Burgess, 2010 Mek1 kinase governs outcomes of meiotic recombination and the checkpoint response. Curr Biol 20: 1707–1716.

Xu, L., M. Ajimura, R. Padmore, C. Klein and N. Kleckner, 1995 NDT80, a meiosis-specific gene required for exit from pachytene in Saccharomyces cerevisiae. Mol Cell Biol 15: 6572–6581.

Yamada, M., N. Hayatsu, A. Matsuura and F. Ishikawa, 1998 Y’-Help1, a DNA helicase encoded by the yeast subtelomeric Y’ element, is induced in survivors defective for telomerase. J Biol Chem 273: 33360–33366.

Zhang, L., K. P. Kim, N. E. Kleckner and A. Storlazzi, 2011 Meiotic double-strand breaks occur once per pair of (sister) chromatids and, via Mec1/ATR and Tel1/ATM, once per quartet of chromatids. Proceedings of the National Academy of Sciences 108: 20036–20041.

Zhao, Q., Y. Xie, Y. Zheng, S. Jiang, W. Liu, et al., 2014 GPS-SUMO: a tool for the prediction of sumoylation sites and SUMO-interaction motifs. Nucleic Acids Res 42: W325–330.

Zickler, D., and N. Kleckner, 1998 The leptotene-zygotene transition of meiosis. Annu Rev Genet 32: 619–697.

Zickler, D., and N. Kleckner, 1999 Meiotic chromosomes: integrating structure and function. Annu Rev Genet 33: 603–754.

Zickler, D., and N. Kleckner, 2015 Recombination, Pairing, and Synapsis of Homologs during Meiosis. Cold Spring Harb Perspect Biol 7.

Zickler, D., and N. Kleckner, 2016 A few of our favorite things: Pairing, the bouquet, crossover interference and evolution of meiosis, pp. 135–148 in Seminars in cell & developmental biology. Elsevier.

